# Selection of potent biologic antagonists of the cannabinoid GPCR CB_2_R from a constrained peptide library

**DOI:** 10.64898/2026.06.29.735442

**Authors:** Robert S Leddy, Ajay Pal, Jamie Plant, Cian McBrien, Ye Li, Hannah Phelan, Sara Linse, Calen Steiner, Colm Collins, David J O’Connell

**Affiliations:** School of Biomolecular & Biomedical Science, University College Dublin, Belfield, Dublin 4, D04V1W8, Ireland; Mucosal Inflammation Program, University of Colorado Hospital, Anschutz Medical Campus, Aurora, CO 80045; Biochemistry and Structural Biology, Lund University, 221 00 Lund, Sweden

## Abstract

Dysregulated gut homing of leukocytes drives chronic inflammation in Crohn’s disease (CD). We employed phage display selection campaigns with libraries of stabilized, constrained peptides against endogenous conformation states of the cannabinoid receptor CB_2_R on human T cells, to discover novel receptor antagonists with potential to inhibit gut homing. Cluster and frequency analysis of 50,000 enriched sequences resulted in expression and functional characterisation of 10 protein candidates using assays of glucose uptake, ERK phosphorylation (pERK) and β-arrestin recruitment. Each candidate antagonised CB_2_R activity with recorded IC50 values of between 5-10 nM. Cannabinoid receptor nanodisc binding experiments and FSEC confirmed CB_2_R selectivity. SLKC_09 with an IC50 of 5.4 nM, was studied in a mouse model of chronic ileitis where it significantly inhibited gut homing of CD4^+^ & CD8^+^ naïve, effector and memory cell types. Our findings highlight an alternative route to therapeutic inhibition of leukocyte trafficking in CD with a biologic inhibitor of CB_2_R.

## Main text

Crohn’s disease (CD) is one of the major types of inflammatory bowel disease (IBD), manifesting as chronic, refractory intestinal disease marked by transmural inflammation and granulomas [1, 2]. The global prevalence of the disease continues to rise with an accelerating incidence in newly industrialized countries and a compounding prevalence in developed ones [3–6]. Standard of care biologic therapies including the anti-TNF antibody Infliximab and the anti-α4β7 antibody Vedolizumab, while demonstrating significant clinical benefit in individual patients, are seen to plateau in a therapeutic ceiling in patient populations. Infliximab benefits approximately 50% of biologic-naïve patients, with up to 50% of responders becoming refractory to the treatment within a year [7]. Vedolizumab improves remission rates in approximately 40% of patients [8–10]. These limitations with IBD therapeutic interventions and the absence of a definitive treatment for CD highlights a clear unmet clinical need in the development of a wider range of therapeutic approaches that could be utilised in a polypharmacology regimen to better manage the disease in more patients.

Self-medication with *cannabis* has been employed by some patients who have reported improvements in levels of abdominal pain, abdominal cramping, diarrhoea, and joint pain [11–13]. Conversely, *cannabis* use for longer than six months is associated with worsened disease outcomes in patients, including a fivefold increase in the risk of surgery [14]. The cannabinoid receptor CB_2_R is a peripherally expressed G protein-coupled receptor (GPCR), predominantly associated with the immune system widely expressed on leukocytes, including T cells that primarily depend on CB_2_R for canonical cannabinoid signalling [14, 15]. Recent evidence describes a role for CB_2_R signalling in promoting the induction of retinoic acid-mediated α4β7 expression on T cells [16]. Taken together, determining whether blockade of the receptor may offer a novel therapeutic avenue for the growing number of IBD patients is of potential significance.

Here we employed a phage display library with a constrained seven-residue variable loop peptide insertion between the two EF-hands of the small and tightly folded calcium binding protein S100G [17], to identify selective antagonists of CB_2_R expressed on human T cells. Lead candidates from computational analysis of deep sequencing from selection rounds were synthesised and assayed for receptor modulation using cellular assays prior to assessing the impact on leukocyte trafficking to the gut in a mouse model of chronic ileitis.

## Results

### Cell surface selection against CB_2_R in multiple cell lines generates a pool of specific peptide binders

Utilising WT Jurkat T cells, which endogenously express CB_2_R, and genetically modified overexpressing *CNR2* CRISPRa HEK293T cells, we established a selection protocol that isolated a pool of enriched CB_2_R-specific peptide binders. In Round 1, WT Jurkat T cells were used, an immortalised human lymphoma cell line which lack CB_1_R expression, exclusively relying on CB_2_R for canonical cannabinoid signalling, therefore mitigating any concerns regarding cannabinoid receptor specificity. Following the completion of Round 1, the number of colonies formed was approximately 56,000 CFU/ml. Round 2 incorporated the incubation of our Round 1 phage stock with *CNR2* CRISPRa HEK293T cells, genetically modified to overexpress CB_2_R, allowing for the amplification of our CB_2_R-specific peptide binders, indicated by an increased CFU/ml of approximately 800,000. Similarly, Round 3 incorporated the incubation of our Round 2 phage stock with WT Jurkat T cells again, further enriching our binders. Given the expression profile of cannabinoid receptors in Jurkat T cells, this Round 3 enrichment ensured we were generating a pool of enriched CB_2_R-specific peptide binders, again indicated by an increased CFU/ml of approximately 1.53 million/ml **(Fig. S1)**. Following next generation sequencing of Rounds 2 and 3 phage stocks, 79,482 7mer peptide binders targeting CB_2_R were identified, 48,969 of which were classified as being CB_2_R-specific **(Table 1)**. Using our computational-based platform, these 48,969 CB_2_R-specific peptides were ranked based on their respective cluster size (i.e. the number of structurally similar peptides) and frequency (i.e. the number of times the peptides is identified in the sequencing data) in order to generate a list of the top 10 CB_2_R-specific peptides. For scientific rigour purposes, a frequency cut-off of > 5 was applied to remove any singleton peptides. Once the top 10 CB_2_R-specific peptides were identified via these criteria, we were able to structurally align them and identify any structural similarities present **(Fig. 1)**. Taken together, these findings highlight the robust nature of phage-based cell surface selection and the importance of a substantial selection outline for the identification of receptor-specific peptide binders. Once identified, these peptide binders can then be ranked accordingly using our computational-based platform and structurally aligned in order to classify any structural similarities among binders prior to protein expression and purification for functional analysis.

**Figure 1:**
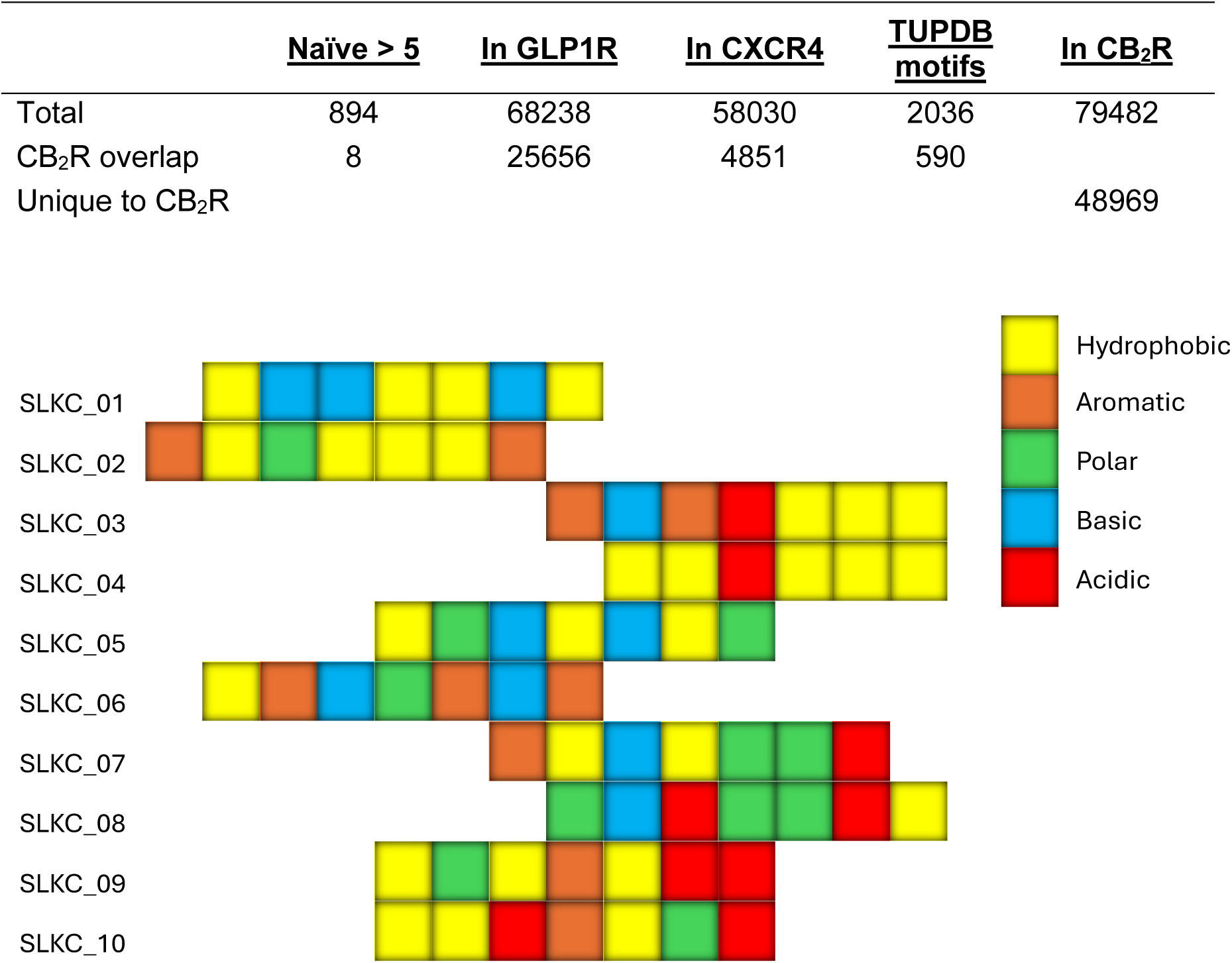
NGS identifies *CNR2*-specific peptide binders. Sequence characteristics for each of the top 10 peptide sequences specific for CB_2_R identified following NGS. These peptides were ranked based on their respective cluster size and frequency, when a frequency cut-off of > 5 was applied.

**Supplementary Figure S1:**
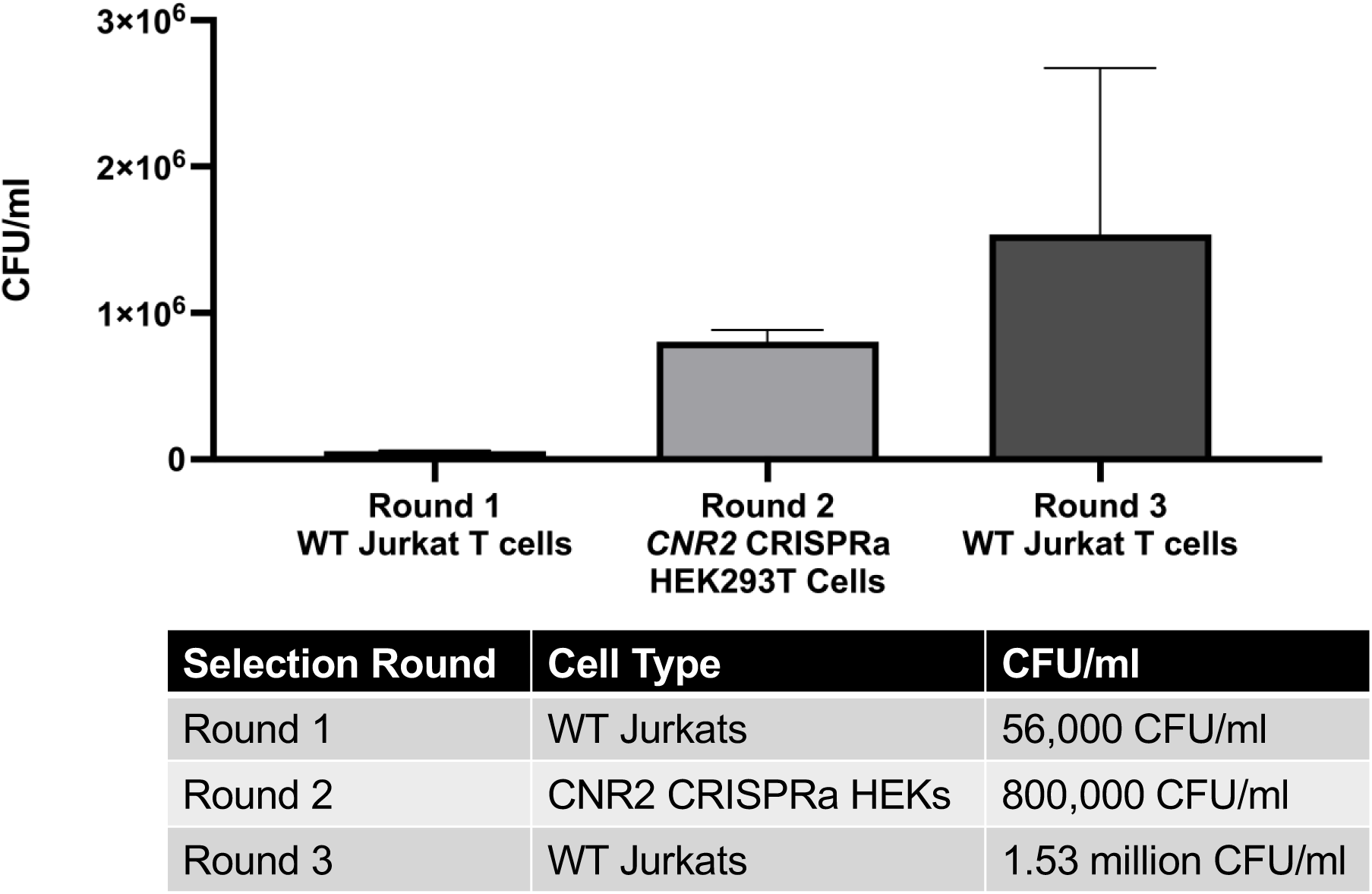
Selection outline utilised for cell surface selection against *CNR2* uses endogenously-expressing and genetically modified overexpressing cell lines for efficient peptide binder amplification. Bar chart and table depicting the CFU/ml values as selection rounds alternate between endogenously *CNR2*-expressing WT Jurkat T cells and genetically modified *CNR2* overexpressing HEK293T cells.

### CB_2_R-specific peptides function as receptor antagonists, inhibiting CB_2_R-mediated glucose uptake into Jurkat T cells

Preliminary unpublished data has described CB_2_R signalling to have a significant role in regulating T cell glucose metabolism, particularly in driving the initial uptake of glucose into T cell following receptor activation as shown via the use of a fluorescent glucose analogue uptake assay utilising the fluorescent analogue 2-NBDG. In order to assess to what extent the identified CB_2_R-specific peptides operated at CB_2_R, we performed this fluorescent glucose analogue uptake assay as a method of functional readout. Jurkat T cells were simultaneously co-treated with 1 μM of the synthetic CB_2_R agonist JWH-133 and each of the top 10 CB_2_R-specific peptides at varying concentrations for 3 h. Vehicle-treated cells received 100% DMSO diluted in the same manner as JWH-133 in glucose-free media in place of any cannabinoid treatment or peptides, whereas a JWH-133-only treatment group in the absence of any peptides was used as a positive control. Following treatment time, the level of fluorescent glucose analogue 2-NBDG take up was quantified via fluorescence detection at Ex/Em of 485nm/520nm. At 3 h, CB_2_R activation in the positive control cohort significantly increased the rate of glucose uptake into treated T cells relative to the vehicle-treated cohort, something that was significantly inhibited by all of the CB_2_R-specific peptides at concentrations of 1 μM, 500 nM, and 100 nM **(Fig. 2)**. In summary, CB_2_R activation drives glucose uptake into T cells which was inhibited significantly by our CB_2_R-specific selektides, indicating that these peptides are operating as receptor antagonists, even at the lower concentration of 100 nM.

**Figure 2:**
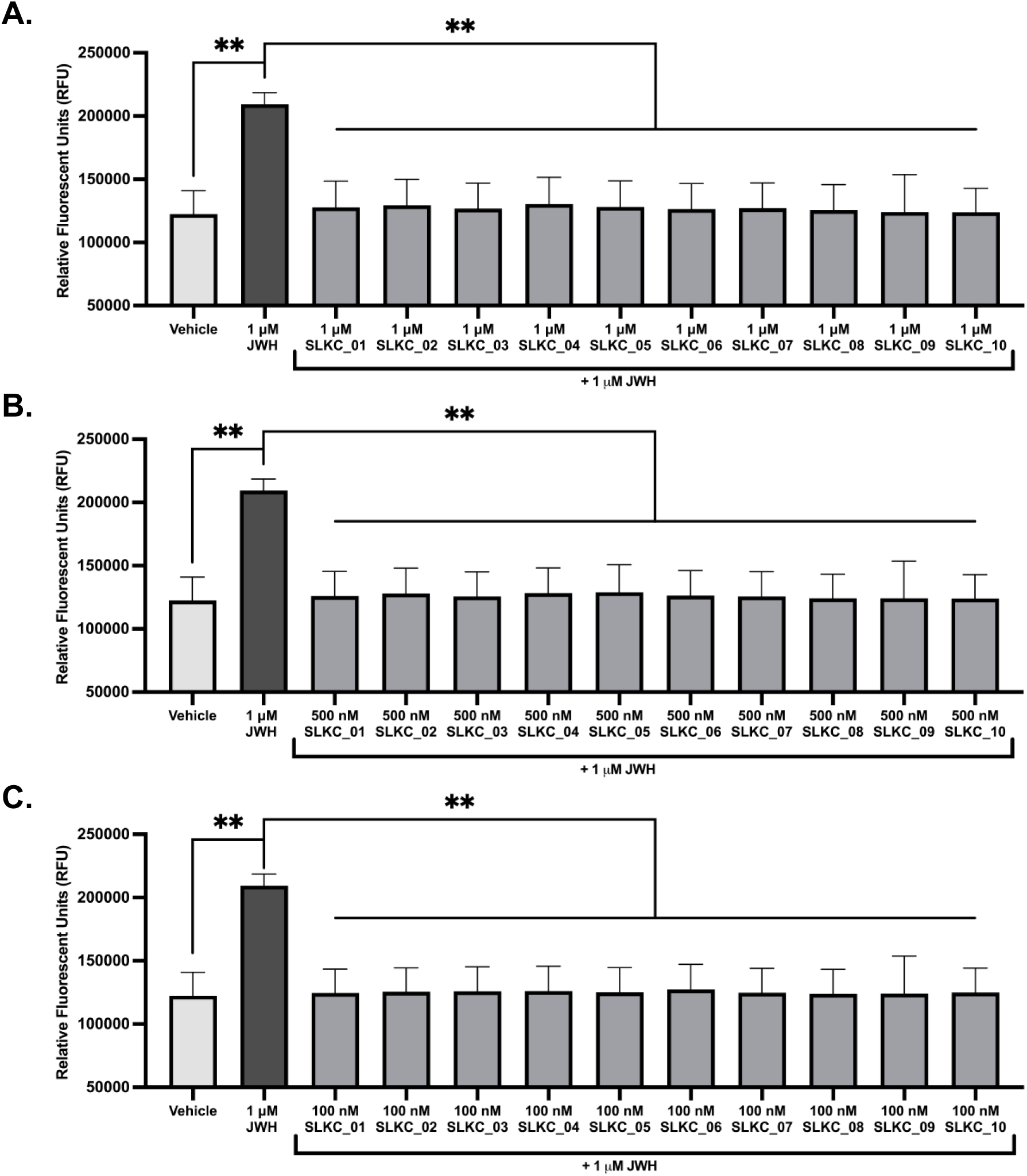
CB_2_R-specific selektides identified following NGS act as receptor antagonists, blocking receptor activation and inhibiting the rapid increase in glucose into T cells. Fluorescent glucose analogue uptake assay into WT Jurkat T cells following 3 h activation of CB_2_R signalling with the synthetic agonist JWH, alongside **(A)** 1 μM **(B)** 500 nM **(C)** 100 nM of the CB_2_R-specific selektides. Bars represent the mean ± SEM of n = 3 independent experiments with n = 3 technical replicates. Statistical significance was determined by unpaired T test comparing each condition to the JWH-133-stimulated control. ** p < 0.01.

### High affinity CB_2_R-specific binding peptides have IC50 values in the low nanomolar range and operate as functional CB_2_R receptor antagonists in a concentration-dependent manner

The fluorescent glucose uptake assay utilised in this study as a functional output of CB_2_R inhibition facilitated the calculation of IC50 values for each of the top 10 CB_2_R-specific peptide binders. Given it had the same inhibitory effect as 1 μM and 500 nM, the 100 nM concentration was used as the starting point in this assay as a means of identifying the lowest inhibitory concentrations for each selektide. The selektides with the lowest IC50 values, inhibiting CB_2_R activation at the lowest nanomolar concentration, and therefore operating with the highest receptor affinity were SLKC_01, SLKC_02, and SLKC_09 with IC50 values in the low nanomolar range of 6.7 nM, 6.5 nM, and 5.4 nM respectively **(Fig. 3D)**. These CB_2_R peptide binders were identified as being potent functional CB_2_R antagonists, inhibiting receptor activation and subsequent downstream CB_2_R-dependent glucose uptake into T cells in a concentration-dependent manner. This functional inhibition was apparent in the nanomolar range from 100 nM to 6 nM, but was completely abolished when concentrations decreased further to 3 nM and 1 nM, indicating loss of their antagonistic features between 6 nM and 3 nM **(Fig 3)**. The S100G scaffold used as a negative control in this assay showed no alteration in the rate of glucose uptake relative to the cohort of cells treated with JWH-133 alone. In summary, our CB_2_R peptide binders operate as potent antagonists of CB_2_R, inhibiting functional output of CB_2_R activation in a concentration-dependent manner with IC50 values in the low nanomolar range.

**Figure 3:**
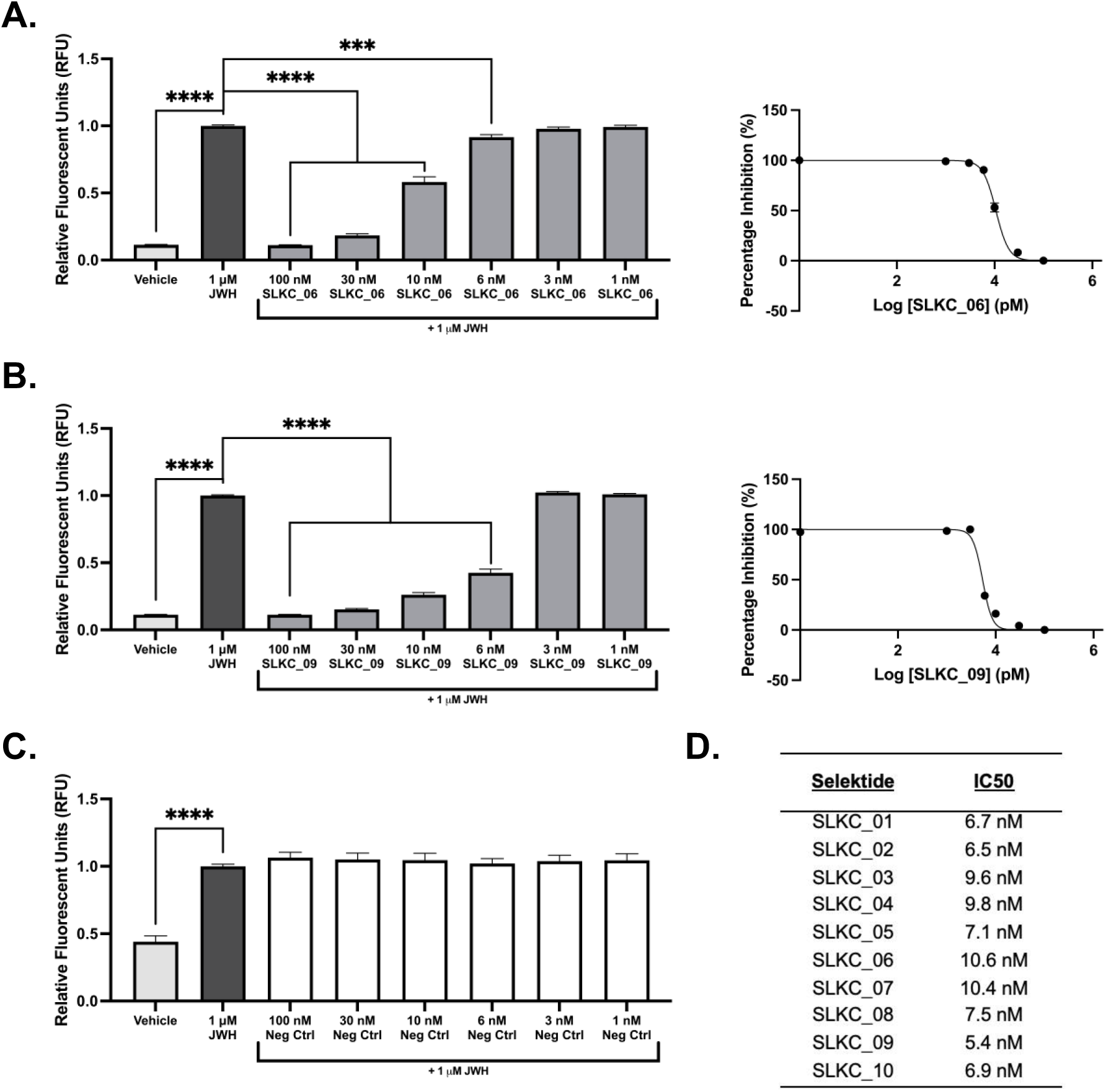
CB_2_R-targeting selektides operate as CB_2_R antagonists, inhibiting the activation of CB_2_R in a concentration-dependent manner and losing inhibitory responses between 6 nM and 3 nM concentrations. Bar charts and XY dot plots highlighting the concentration-dependent inhibitory effect of the **(A)** most and **(B)** the least potent CB_2_R-targeting selektides, in addition to **(C)** the S100G negative control on receptor activation in a fluorescent glucose analogue uptake assay into WT Jurkat T cells following 3 h activation of CB_2_R signalling. **(D)** Table displaying the IC50 values calculated for each tested selektide in the fluorescent glucose analogue uptake assay. Bars represent the mean ± SEM of n = 3 independent experiments with n = 3 technical replicates. Statistical significance was determined by unpaired T test comparing each condition to the JWH-133-stimulated control. *** p < 0.001; **** p < 0.0001. Colour schematics depict the structural characteristics of selected peptides.

**Figure 4:**
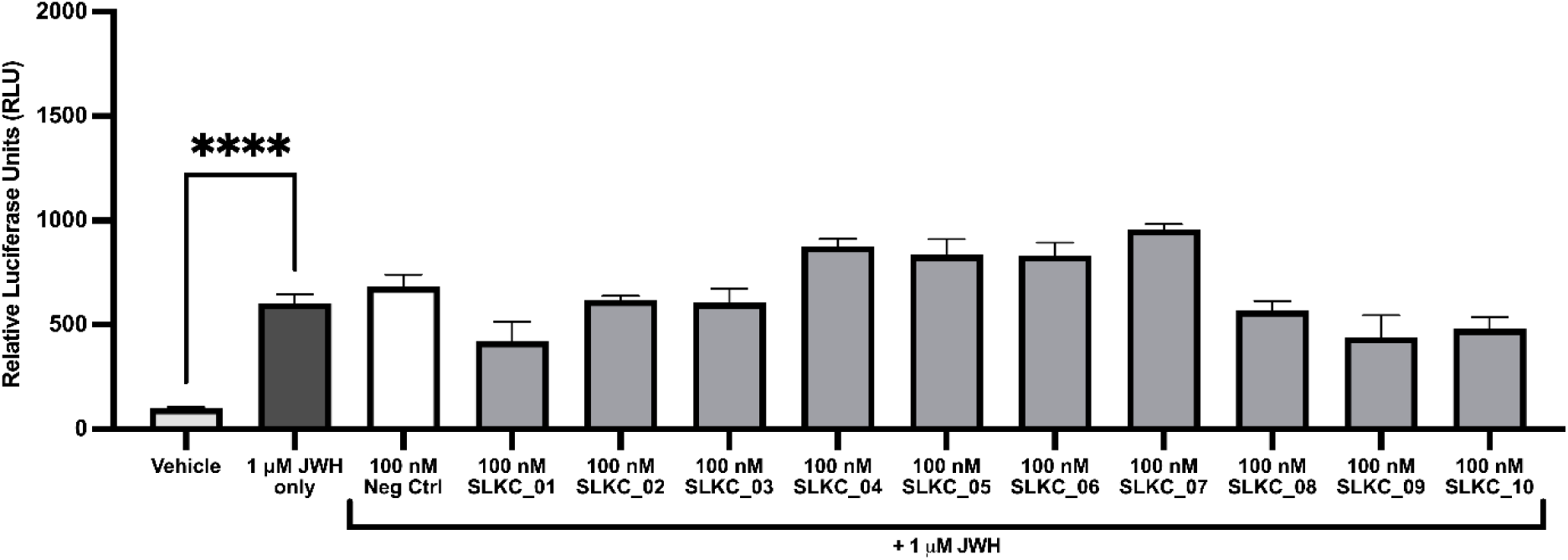
The top 10 highest affinity CB_2_R-targeting selektides show distinct profiles in their modulation of intracellular G-protein independent β-arrestin recruitment. Bar chart highlighting the rate of β-arrestin recruitment to the activated receptor in CB_2_R-Tango transfected HTLA cells when co-administered with both 1 μM JWH-133 and 100 nM of respective selektide for 24 h. Bars represent the mean ± SEM of n = 3 independent experiments with n = 3 technical replicates. Statistical significance was determined by unpaired T test comparing each condition to the JWH-133-stimulated control. **** p < 0.0001.

### High affinity CB_2_R-specific selektides operate as receptor antagonists but show distinct profiles in modulating intracellular β-arrestin recruitment to CB_2_R

Using the β-arrestin recruitment assay presented herein, the top 10 CB_2_R-targeting selektides were screened against CB_2_R for modulation of G-protein independent β-arrestin recruitment, with quantitative measurements subsequently determined via luminescence readings. HTLA cells transfected with the CB_2_R-Tango plasmid were simultaneously co-treated with 1 μM of the synthetic CB_2_R agonist JWH-133 and our selektides at 100 nM for 24 h. The working concentration of 100 nM was chosen based off the previously described fluorescent glucose analogue uptake assay. Vehicle-treated cells received 100% DMSO diluted in the same manner as JWH-133 in supplemented DMEM media in the place of any cannabinoid treatment or selektides, whereas a JWH-only treatment group was used as a positive control. The screened selektides displayed trends towards distinct profiles in their modulation of β-arrestin recruitment to the activated CB_2_R. CB_2_R activation in the positive control cohort significantly increased the rate of β-arrestin recruitment in transfected cells in comparison to vehicle-treated cohort, in a manner that was expected given JWH-133 is well-documented to be a highly specific CB_2_R agonist [18, 19]. SLKC_01, SLKC_09, and SLKC_10 all displayed a trend towards the inhibition of agonist-induced β-arrestin recruitment, whereas SLKC_04, SLKC_05, SLKC_06, and SLKC_07 looked to increase β-arrestin recruitment when administered alongside the CB_2_R agonist. Additionally, SLKC_02, SLKC_03, and SLKC_08 showed no change in the recruitment of β-arrestin when compared with the JWH-133 alone cohort of treated cells. The S100G scaffold used as a negative control in this assay showed no alteration in the rate of β-arrestin recruitment **(Fig. 5)**. Despite these alterations being deemed statistically insignificant, there are clear trends in the data that appear to indicate distinct profiles of these CB_2_R-targeting selektides in their modulation of β-arrestin recruitment which may be due to distinctions in their sequence makeup and resulting signalling bias.

**Figure 5:**
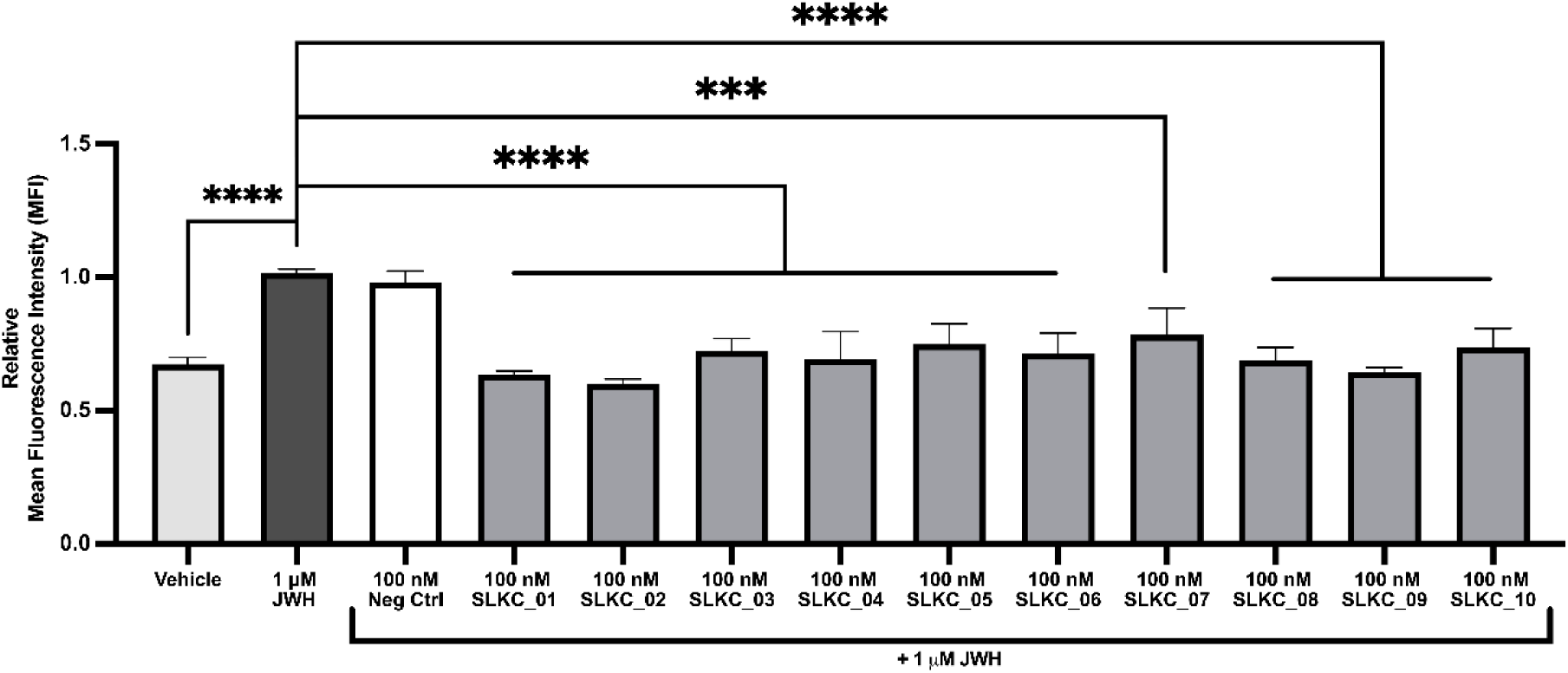
Selektide-mediated attenuation of CB_2_R-induced ERK phosphorylation assessed by flow cytometry. Intracellular pERK levels were measured by flow cytometry in cells stimulated with the CB_2_R agonist JWH-133 (1 µM) in the presence or absence of selektide SLKC_01–SLKC_10 (100 nM each). Data is expressed as relative mean fluorescence intensity (MFI), normalised to the 1 µM JWH-133 positive control. A 100 nM negative control peptide co-treated with JWH-133 was included to control for non-specific peptide effects. Bars represent the mean ± SEM of n = 3 independent experiments with n = 3 technical replicates. Statistical significance was determined by unpaired T test comparing each condition to the JWH-133-stimulated control. *** p < 0.001; **** p < 0.0001.

### Selektide-mediated attenuation of ERK phosphorylation

To investigate the capacity of the selektides to modulate CB_2_R-mediated ERK signalling, intracellular pERK levels were quantified by flow cytometry following treatment with the selektide candidates (SLKC_01-SLKC_10) at 100 nM in the presence of the CB_2_R agonist JWH-133 (1 µM). Relative mean fluorescence intensity (MFI) was used as a measure of ERK phosphorylation across all conditions. Stimulation with 1 µM JWH-133 alone produced a significant increase in pERK relative to vehicle-treated cells, confirming robust agonist-induced ERK activation under these experimental conditions. The 100 nM negative control peptide co-administered with JWH-133 did not alter pERK levels compared to agonist stimulation alone, demonstrating that non-specific peptide effects did not account for any observed attenuation. Co-treatment with selektides SLKC_01, SLKC_02, and SLKC_09 resulted in a statistically significant reduction in pERK relative to the JWH-133-stimulated positive control, with these candidates reducing relative MFI values to approximately 0.60– 0.65 of the positive control, suggesting meaningful antagonism of agonist-induced ERK activation. The remaining selektides also reduced pERK levels relative to the JWH-133 control, with relative MFI values ranging from 0.66 to 0.80. Collectively, these data indicate these selektides are capable of attenuating CB_2_R-mediated ERK phosphorylation, with SLKC_01, SLKC_02, and SLKC_09 emerging as the most potent candidates at the concentration tested **(Fig. 5)**.

### SLKC_09 and SLKC_09_ABD reduce CD4⁺ T cell numbers across gut-associated lymphoid compartments

To evaluate the effects of selektide SLKC_09 and its modified variant with an albumin-binding domain SLKC_09_ABD on T cell trafficking in vivo, total T cell numbers were quantified across multiple phenotypically distinct subpopulations in the mesenteric lymph node (MLN) and intestinal lamina propria (LP) following selektide treatment relative to PBS vehicle control. Both SLKC_09_ABD and SLKC_09 significantly reduced total CD4⁺ T cell numbers compared to PBS-treated animals in the MLN, while in the LP only the unmodified SLKC_09 significantly reduced total CD4⁺ T cells infiltration **(Fig. 6A)**. Analysis of the effector CD44^High^CD62L^Low^ T cell subtype revealed significant reductions in the MLN for both SLKC_09 and SLKC_09_ABD, with SLKC_09 also significantly reducing the LP counterpart **(Fig. 6B)**. Within the central memory CD4⁺CD44^High^CD62L^High^ cohort, both selektides significantly reduced the numbers of these cells in the MLN, though neither produced a statistically significant reduction in the LP for this subset **(Fig. 6C)**. For the naïve CD4⁺CD44^Low^CD62L^High^ population, SLKC_09 significantly reduced LP cell numbers, while both SLKC_09 and the modified SLKC_09_ABD resulted in significant reduction in the corresponding MLN population **(Fig. 6D)**.

**Figure 6:**
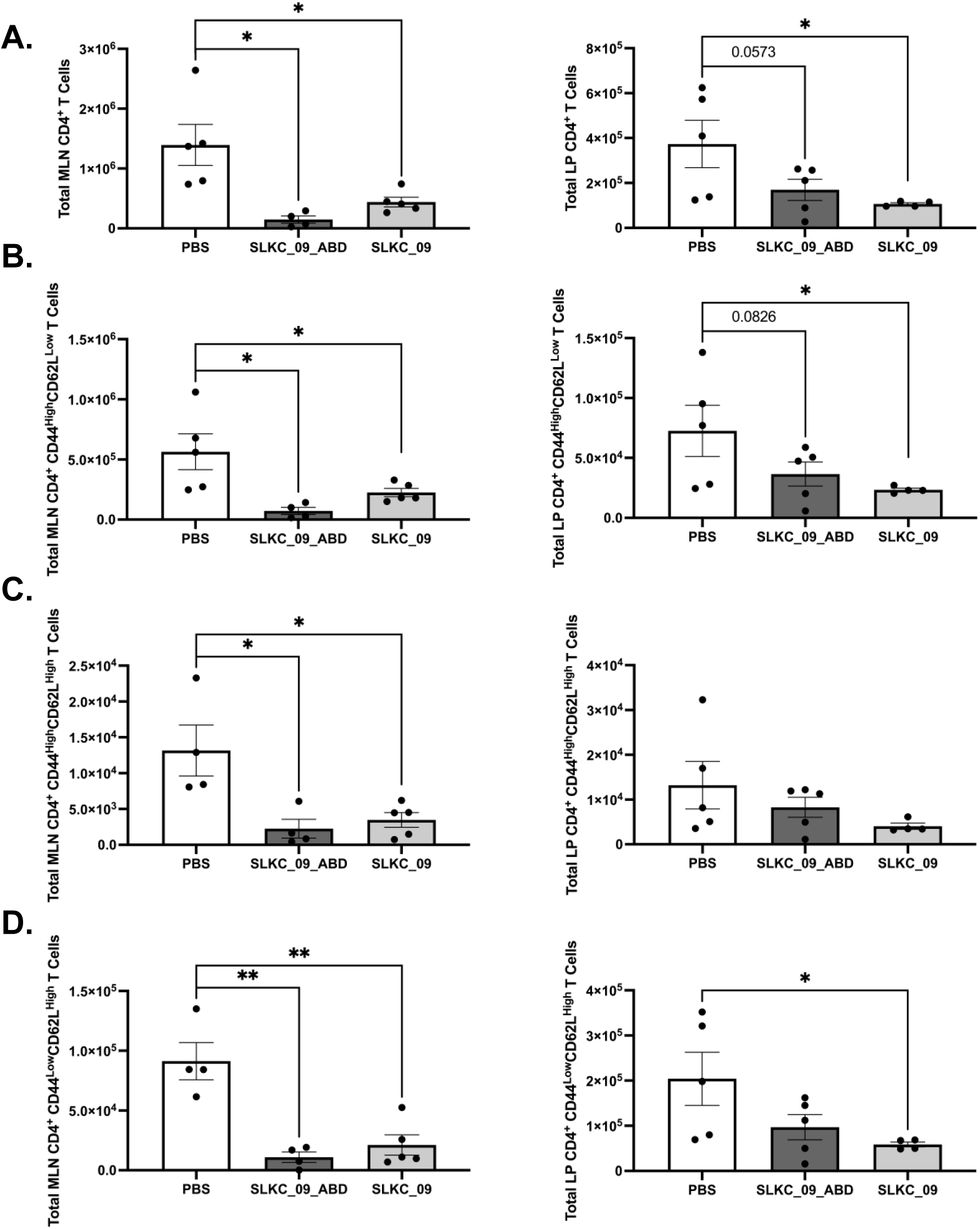
SLKC_09 and SLKC_09_ABD reduce CD4⁺ T cell accumulation across phenotypically distinct subpopulations in the mesenteric lymph node and intestinal lamina propria. Total CD4⁺ T cell numbers were quantified by flow cytometry across four phenotypically defined subpopulations in the mesenteric lymph node (MLN) and intestinal lamina propria (LP) following treatment of TNF^ΔARE^ mice with SLKC_09_ABD or SLKC_09 relative to PBS vehicle control. **(A)** Total CD4⁺ T cells in the MLN and LP. **(B)** Effector CD4⁺CD44^High^CD62L^Low^ T cells in the MLN and LP. **(C)** Central memory CD4⁺CD44^High^CD62L^High^ T cells in the MLN and LP. **(D)** Naïve CD4⁺CD44^Low^CD62L^High^ T cells in the LP and MLN. Bars represent mean ± SEM of n = 5 animals per group. Statistical comparisons were made relative to PBS control. * p < 0.05; ** p < 0.01; p values shown where differences trended toward significance.

**Supplementary Figure S2:**
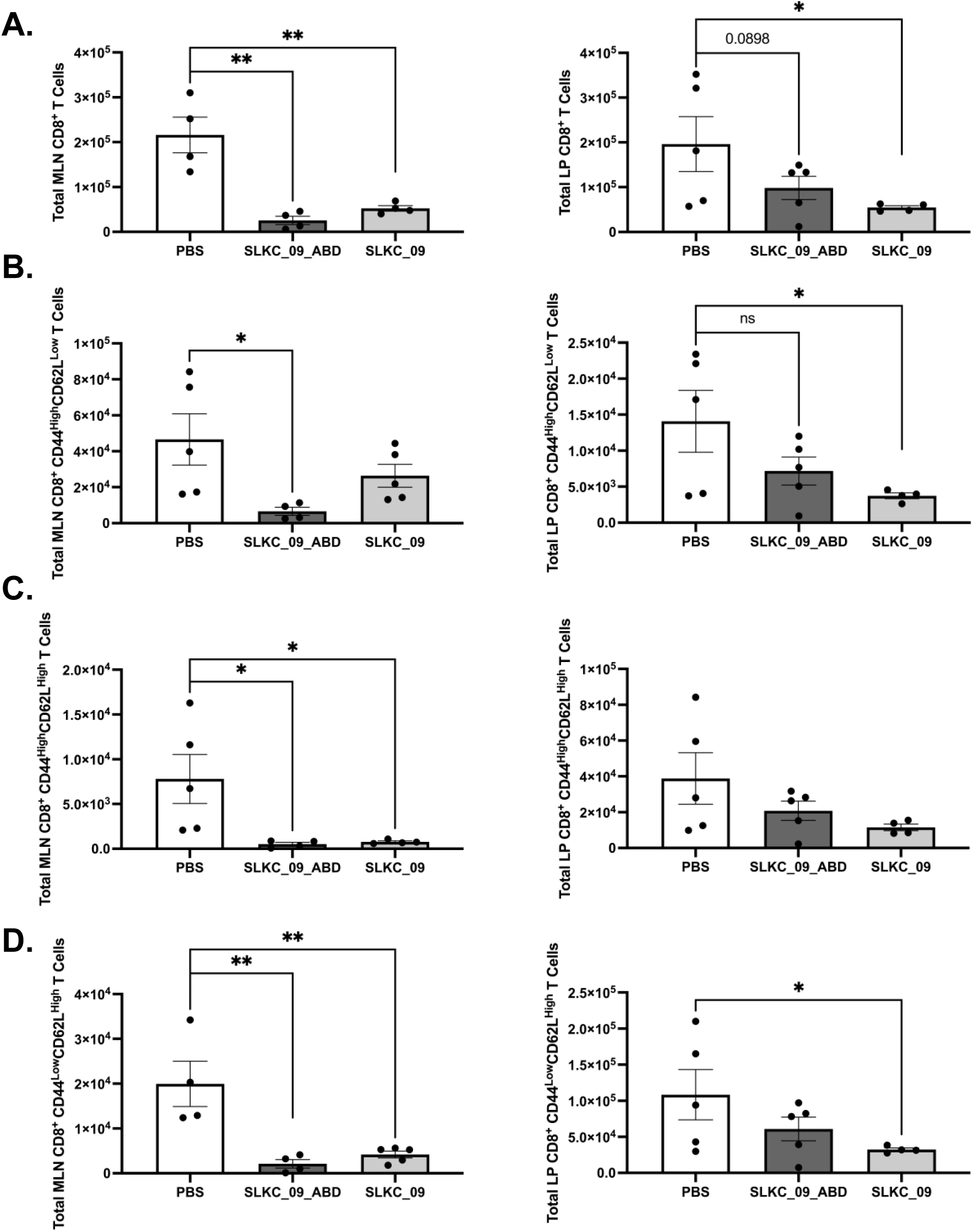
SLKC_09 and SLKC_09_ABD broadly suppress CD8⁺ T cell subsets across gut-associated lymphoid compartments. Total CD8⁺ T cell numbers were quantified by flow cytometry across four phenotypically defined subpopulations in the mesenteric lymph node (MLN) and intestinal lamina propria (LP) following treatment of TNF^ΔARE^ mice with SLKC_09_ABD or SLKC_09 relative to PBS vehicle control. **(A)** Total CD8⁺ T cells in the MLN and LP. **(B)** Effector CD8⁺CD44^High^CD62L^Low^ T cells in the MLN and LP. **(C)** Central memory CD8⁺CD44^High^CD62L^High^ T cells in the MLN and LP. **(D)** Naïve CD8⁺CD44^Low^CD62L^High^ T cells in the LP and MLN. Bars represent mean ± SEM of n = 5 animals per group. Statistical comparisons were made relative to PBS control. * p < 0.05; ** p < 0.01; p values shown where differences trended toward significance.

Taken together, this data demonstrates that both SLKC_09 and SLKC_09_ABD broadly suppress the accumulation of CD4⁺ T cell subsets across multiple gut-associated lymphoid compartments in the MLN and LP, with consistent and reproducible effects observed across naïve, effector memory, and central memory populations.

### SLKC_09 and SLKC_09_ABD reduce gut-homing α4β7⁺ T cell numbers in the lamina propria and mesenteric lymph nodes via the inhibition of α4β7 expression

To determine whether selektide-mediated suppression extended to gut-homing integrin-expressing T cell populations, total α4β7⁺ cell numbers were quantified within both CD4⁺ and CD8⁺ compartments in the LP and MLN following treatment with SLKC_09 or SLKC_09_ABD relative to PBS vehicle control.

SLKC_09 significantly reduced LP CD4⁺α4β7⁺ T cell numbers compared to PBS-treated animals, while SLKC_09_ABD trended toward reduction without reaching significance. Conversely, in the MLN SLKC_09_ABD produced a significant reduction in CD4⁺α4β7⁺ T cell numbers, while the reduction observed with SLKC_09 alone did not achieve statistical significance in this compartment **(Fig. 8A)**. For the CD8⁺α4β7⁺ gut-homing population, neither selektide resulted in a significant alteration in the total LP numbers, whereas both SLKC_09_ABD and SLKC_09 significantly reduced MLN CD8⁺α4β7⁺ T cell numbers **(Fig. 8B)**.

**Figure 8:**
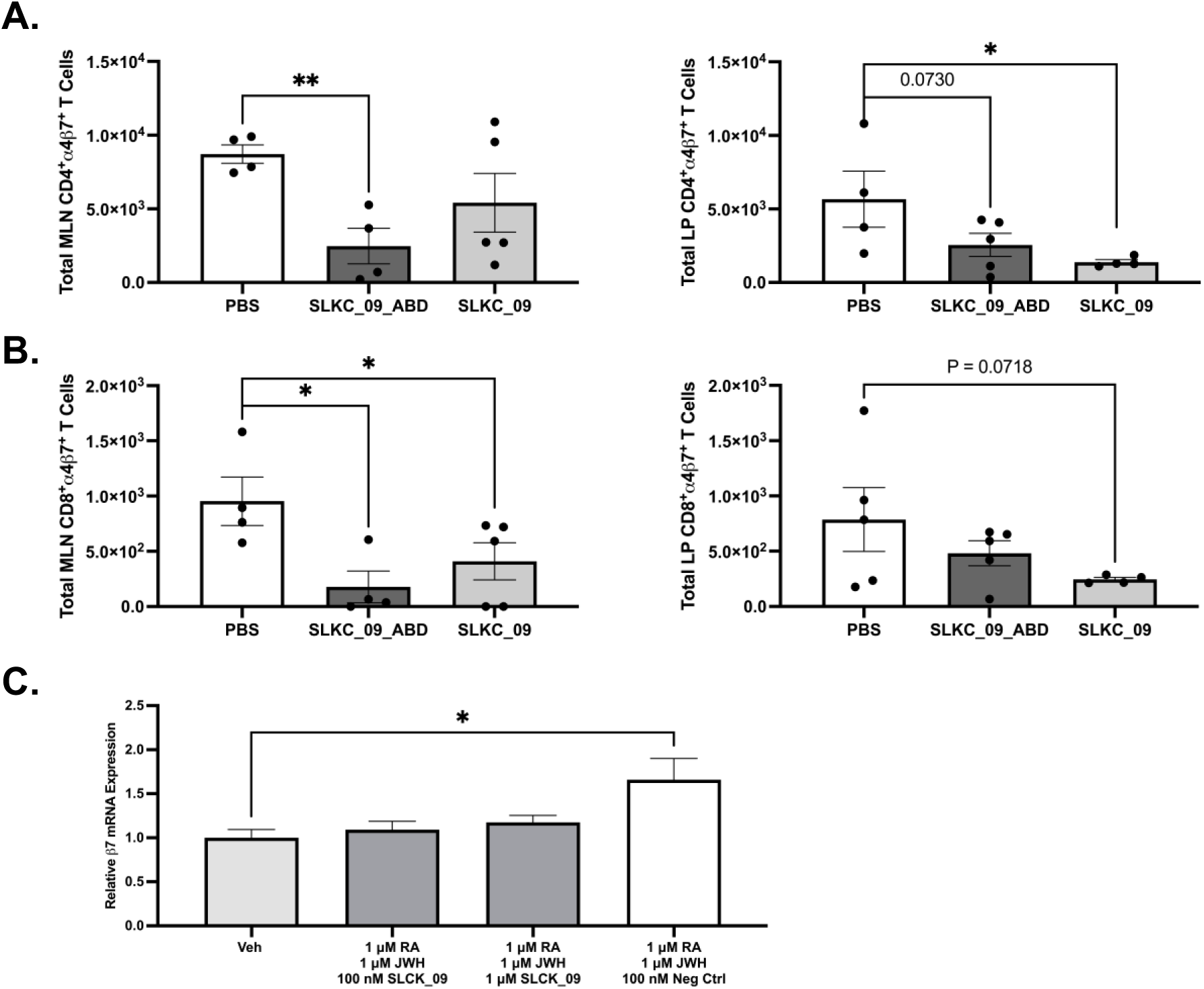
SLKC_09 decreases gut-homing α4β7⁺ CD4⁺ and CD8⁺ T cell numbers in the lamina propria and mesenteric lymph node via the inhibition of α4β7 expression on T cells. Total numbers of gut-homing α4β7-integrin-expressing T cells were quantified by flow cytometry in the LP and MLN of TNF^ΔARE^ mice following treatment with SLKC_09_ABD or SLKC_09 relative to PBS vehicle control. **(A)** LP and MLN CD4⁺α4β7⁺ T cells**. (B)** LP and MLN CD8⁺α4β7⁺ T cells. Bars represent mean ± SEM of n = 5 animals per group. Statistical comparisons were made relative to PBS control. * p < 0.05; ** p < 0.01; p values shown where differences trended toward significance. **(C)** Real time qPCR shows SLKC_09 to inhibit the CB_2_R-driven induction of α4β7 expression on T cells at an mRNA level. Bars represent the mean ± SEM of n = 3 independent experiments with n = 3 technical replicates. Statistical significance was determined by unpaired T test comparing each condition to the JWH-133-stimulated control. * p < 0.05.

To investigate the mechanism through which SLKC_09 could potentially decrease the accumulation of α4β7-expressing T cells in the MLN and LP of TNF^ΔARE^ mice, RT-qPCR was used to assess the expression of integrin β7 in Jurkat T cells following exposure to the CB_2_R agonist JWH-133 and SLKC_09, which was found to inhibit the induction of CB_2_R-driven β7 expression at an mRNA level at both concentrations tested.

This data indicates that SLKC_09 suppresses the accumulation of gut-homing α4β7-expressing T cells in a compartment- and lineage-dependent manner, with particularly pronounced and consistent effects observed in the MLN across both CD4⁺ and CD8⁺ populations, via the inhibition of CB_2_R-driven α4β7 expression on T cells.

## Discussion

With rising global incidence and the incurable and chronic nature of CD & UC, IBD is one of the most prevalent and costly gastrointestinal disorders worldwide. According to a report conducted by Berkshire Hathaway in April 2025, the market for IBD therapeutics is experiencing a period of significant growth, projected to reach an unprecedented high of $33.99 billion by 2033, underlining the growing number of IBD patients and the growing therapeutic need. Globally, *cannabis* and *cannabis*-based products, known as cannabinomimetics, have grown increasingly popular among IBD patients as a complementary therapeutic aid, acting as an opioid alternative in providing chronic pain relief, managing and suppressing overactive gastrointestinal motility, and improving appetite. Numerous questionnaire-based studies and small clinical trials have indicated that self-medication for relief from IBD-associated symptoms are the main reasons for *cannabis* use in IBD patients [11–13]. A study with over two million IBD patients in 2015 recorded a higher incidence and earlier onset of *cannabis* use compared to healthy controls [20], with a survey in a paediatric IBD clinic reporting regular use in 70% of patients with a median age of 18.7 [21].

However, cannabinoids have failed to show any evidence of disease improvement in multiple interventional clinical trials in CD and UC [22]. While *cannabis* may provide temporary symptomatic relief, chronic use for longer than six months is associated with worsened disease outcomes in CD patients, including a five-fold increase in surgical risk, despite reported symptomatic improvements [23]. This disparity between patient-reported symptom relief and the lack of clinical benefit may stem from cannabinoid-associated psychoactive effects, potentially helping patients feel better without improving the underlying disease. A study in a chronic ileitis model, in which affected mice develop progressive and chronic ileitis that mimics many features of CD, found that blockade of CB_2_R signalling attenuated chronic murine ileitis, as demonstrated by improved histological scoring and decreased inflammatory cytokine expression [24]. The tissue expression profile of CB_2_R, reviewed extensively elsewhere, makes CB_2_R all the more interesting as a target in IBD [16, 25–31]. The distinct expression profile of this receptor in IBD, together with the findings that *cannabis* use can lead to disease worsening, suggest that overactive endocannabinoid signalling, may have a crucial role in the development of IBD pathogenesis.

The peripheral nature of CB_2_R on T cells, may offer a novel therapeutic mechanism avoiding the negative psychiatric and psychoactive effects associated with CB_1_R as previously observed with Rimonabant^®^, a selective CB_1_R antagonist, initially approved for the treatment of obesity but withdrawn due to complications including anxiety, depression, and suicidal ideology in patients [32, 33]. This study takes advantage of the discrete expression profile of CB_2_R on Jurkat T cells, that express high levels of functional CB_2_R in a constitutive manner [34, 35], and lack CB_1_R expression [15]. This ensures that, following extensive rounds of phage display-based cell surface selection, an enriched pool of highly selective CB_2_R peptide binders is generated with potential as a novel therapeutic for IBD.

The functional assays utilised in the study herein is a fluorescent glucose analogue uptake assay, employing 2-NBDG, a fluorescently-labelled deoxyglucose analogue, as a probe for the detection of glucose taken up by cultured cells, the subsequent level of which can be quantified via the use of fluorescent filters operating at Ex/Em of 485nm/535nm. Previously conducted unpublished data has indicated a role for CB_2_R signalling in T cell glucose metabolism, particularly in significantly driving the uptake of glucose in to T cells in a CB_2_R-dependent manner following receptor activation. As a direct result, the assay employed in this study operates as a functional readout of CB_2_R activation or inhibition, acting as a high-throughput screen of our generated CB_2_R-targeting selektides and facilitating the deciphering of their pharmacodynamic profile. Following three rounds of affinity selection using our patented library of constrained peptides and subsequent NGS, 10 of the highest ranked selektides were purified as soluble proteins and assessed in this assay in terms of their pharmacodynamic profile at CB_2_R, all of which operated as functional receptor antagonists, significantly inhibiting the CB_2_R-mediated uptake of glucose into Jurkat T cells.

We also utilised a luciferase reporter based Tango system first developed by Barnea *et al* [36] to screen our selektides against CB_2_R for β-arrestin recruitment to the activated receptor. β-arrestin recruitment is an important and classical aspect of GPCR function for the termination of G-protein dependent signalling, inducing G-protein dissociation and ultimately leading to both receptor desensitisation and internalisation [37]. This Tango system involves the introduction of three exogenous genetic elements: a protein fusion consisting of β-arrestin2 with a TEV protease, a tTA tethered to CB_2_R via TEV cleavage sites preceded by a sequence to promote arrestin recruitment, and a luciferase reporter gene whose transcription is triggered upon tTA transcription factor translocation to the nucleus, released upon β-arrestin2 recruitment [37]. This assay offers a selective readout to the target receptor of interest, alongside high sensitivity and robustness due to signal integration, making it an effective candidate for high-throughput ligand screening against specific GPCR targets. In addition to agonist-induced β-arrestin recruitment, this assay can also be utilised to quantify the activity of both antagonists and allosteric modulators. The results herein and the distinctions observed with our selektides in relation to β-arrestin recruitment is in agreement with the concept of ligand bias or functional selectivity, a concept where different ligand structures can elicit distinct receptor signalling cascades at a single receptor. To delve deeper into this concept and to get a greater understanding of the signalling cascades induced by our CB_2_R-targeting selektides, in addition to their proposed antagonism, we also assessed the activation of the G-protein-dependent signalling arm following exposure to our selektides, paying particular attention to phosphorylated ERK, a well-documented downstream effector following CB_2_R activation. While there appear to be differences between the manner in which these selektides modulate agonist-induced β-arrestin recruitment, there is consistency between the tested peptides in the assessment of downstream pERK levels, which all appear to significantly inhibit CB_2_R-agonist induced ERK1/2 phosphorylation, thus antagonising G-protein signalling. Given the inhibition of CB_2_R-driven glucose uptake, together with the reduction in agonist-mediated pERK levels, we can suggest that these selektides are operating as CB_2_R antagonists with high affinities. However, the results generated from the β-arrestin recruitment assay and the distinctions in which these selektides modulate β-arrestin recruitment allows us to stratify these further. SLKC_01, SLKC_09, and SLKC_10 all appear to function as classical neutral antagonists, inhibiting agonist-driven responses for both G-protein-mediated downstream ERK signalling and arrestin recruitment. Contrastingly, selektides SLKC_04, SLKC_05, SLKC_06, and SLKC_07 show characteristics of functional selectivity, inhibiting agonist-induced ERK phosphorylation but still enhancing the recruitment of β-arrestin. These ligands display characteristics not classically associated with a traditional antagonist, instead having the profile of β-arrestin biased agonists. This means the ligand inhibits G-protein-mediated signalling, such as the G_i/o_/ERK pathway while promoting β-arrestin recruitment, potentially via the stabilisation of the receptor in a conformation that is functionally selective towards β-arrestin recruitment over G-protein activation. Functionally, these ligands behave as an antagonist for one signalling branch, G-protein signalling in this case, and a partial agonist for another in β-arrestin recruitment, producing pathway-selective signalling rather than uniform inhibition. A well-known example of this type of ligand is TRV027, an angiotensin II peptide analogue operating at the angiotensin II type 1 receptor studied extensively in heart failure, which blocks classical G_q_-mediated signalling yet enhances the recruitment of β-arrestin1/2 [38]. Alternatively, a ligand that reduces pERK phosphorylation while sparing β-arrestin recruitment, especially when coapplied with an agonist, shows pathway selective antagonism (also known as G-protein-biased antagonism). This is the case with selektides SLKC_02, SLKC_03, and SLKC_08, meaning these selektides selectively block or weaken G-protein-mediated signalling such as the G_i/o_/ERK cascade, yet do not interfere or engage with the events required for β-arrestin recruitment. It may be possible that these peptides are binding an allosteric site, selectively reducing G-protein signalling efficacy without affecting arrestin recruitment. Either way, the functional outcome is preserved arrestin recruitment alongside suppressed G-protein output, effectively creating a bias in signalling and reflecting functional selectivity rather than uniform inhibition. An example of this is the peptide PTH (1-34) which binds to the parathyroid hormone receptor 1 and acts as a biased or pathway-selective negative allosteric modulator that reduces G-protein-driven cAMP and ERK phosphorylation while maintaining regular β-arrestin recruitment elicited by full agonists [39].

As reported by Leddy et al., (2026), CB_2_R signalling is pivotal in promoting the induction of retinoic acid-mediated α4β7 expression on T cells, the heterodimeric integrin responsible for the specific trafficking of T cells to the gut and gut-associated intestinal tissues, contributing massively to the chronic inflammation characteristic of IBD. Additionally, this study also highlighted that the deletion of *CNR2*, the gene encoding CB_2_R, attenuated chronic murine ileitis in 20-week-old TNF^ΔARE/+^ mice, indicating that inhibition of CB_2_R signalling may offer potential therapeutic benefit in IBD. To assess whether our selektides could mimic this by means of CB_2_R antagonism, SLKC_09 and an iteration of this selektide with an albumin binding domain fusion were administered subcutaneously in mice of the same model. Given it appeared to have the highest potency and looked to operate as a typical receptor antagonist in the functional assays herein, SLKC_09 was picked for this in vivo study, which was seen to inhibit the homing of both CD4^+^ and CD8^+^ T cells of naïve, effector, and memory phenotype relative to PBS-treated mice. Using RT-qPCR, we were also able to show that this selektide inhibited the induction of retinoic acid-mediated α4β7 expression on T cells, showing that our CB_2_R-targeting peptide antagonists inhibit the expression of α4β7 and therefore reduce subsequent leukocyte trafficking to gut, potentially being beneficial in the treatment of IBD. CB_2_R agonism has been reported in some colitis models as being beneficial in reducing mucosal inflammation, although conversely the *CNR2*^-/-^ mice studied did not experience aggravated colitis [25]. Our data in a chronic disease model confirms antagonism as an alternative route to therapeutic inhibition of leukocyte trafficking in CD.

Given the lipid-based nature of the endocannabinoid system, the resources available do not indicate that any peptide-based drugs targeting either CB_1_R or CB_2_R are currently on the market, with the majority of clinically used compounds targeting cannabinoid receptors being small molecules [40]. However, some efforts have been made to discover novel cannabinoid receptor peptide binders. An example of this are hemopressin (Hp) and the related peptides VD-Hp and RVD-Hp found in rodents and humans which have been shown to exhibit high- affinity binding to CB_1_R and lower affinities for CB_2_R [41]. Additionally, researchers have designed and characterised bicyclic peptides based on the cyclotide vodo-C1 derived from the sweet violet plant, with their lack of functional activation of CB_2_R and partial radioligand displacement being consistent with weak competitive antagonists [42]. This study highlights our ability to generate highly specific CB_2_R peptide antagonists from constrained peptide library which could offer potential as novel therapeutic strategies for the treatment of IBD, offering several advantages over traditionally used small molecules and antibodies currently dominating the IBD and CB_2_R-targeting markets such as enhanced target specificity, better tissue penetration and cellular internalisation, and their design for various delivery routes including injection, nasal administration, or even topical application [43, 44]. Peptides often bind with higher affinity and specificity to their intended targets due to their size and structure traditionally being similar to those of natural biological molecules [43]. This appears to be the case with our CB_2_R peptide binders, operating as potent functional receptor antagonists in a concentration-dependent manner with IC50 value in the low nanomolar range, the concentration required to inhibit a biological process by 50%. In particular, the highest affinity CB_2_R-binding selektides SLKC_01, SLKC_02, SLKC_09 inhibited receptor activation and subsequent downstream CB_2_R-dependent glucose uptake into Jurkat T cells with IC50 values in the low nanomolar range of 6.7 nM, 6.5 nM, and 5.4 nM, respectively. Generally, the IC50 value of peptides drugs are highly variable. For example, a C1q-binding peptide that inhibits the classic complement pathway showed IC50 values between 2-6 μM [45]. In breast cancer cell lines, some peptides show IC50 values ranging from 16.3 to 137 μM [46], whereas peptides targeting major histocompatibility complexes molecules present with IC50 values in the range of 50-500 nM [47]. Whilst appearing to be highly variable, the IC50 generated for our selektides are in the low nanomolar range and are much lower than these examples, indicating our ability to utilise a constrained peptide library to generate highly potent and specific CB_2_R antagonists. Given the high level of variability within our constrained peptide library used in this study, the antagonists generated also display distinct profiles in terms of their pharmacodynamic profile and functional selectivity, not only functioning as classical neutral antagonists. This is critical and adds significant credence to the studies herein for the generation of potentially therapeutic peptide ligands against GPCRs previously deemed difficult to target, such as CB_2_R. The presence of neutral antagonists, functionally selective antagonists, and negative allosteric modulators in our top 10 CB_2_R-specific selektides reflects the structural and functional diversity captured by our peptide library, allowing the engagement of multiple receptor conformations and intracellular signalling pathways whilst also speaking to the complicated nature of GPCR signalling. The diversity amongst our selektides is potentially therapeutically significant as it enables more precise modulation of CB_2_R activity: classical antagonists provide robust inhibition when needed, whereas allosteric modulators/functionally selective binders can subtly regulate receptor function to preserve beneficial signalling and inhibit signalling that is not. Importantly, while this study is mainly focused on the IBD patient cohort and CB_2_R, such mechanistic variety expands the potential patient populations, as individuals who cannot tolerate full antagonists may benefit from modulators or pathway-biased peptides, opening opportunities to target specific disease-relevant pathways and ultimately increasing the numb er of indications that could be treated safely and effectively.

## Conclusion

We have shown in this study that the use of a patented, constrained peptide library based on a small and highly stable Ca^2+^-binding protein scaffold, combined with phage-based cell surface selection against CB_2_R, enables the identification of highly specific peptide binders. Notably, we were able to identify a ranked list of binders from this library that are highly specific to CB_2_R which operate as potent functional CB_2_R antagonists, operating as either as classical antagonist or antagonists with distinct degrees of functional selectivity. The identification of a peptide antagonists specific to CB_2_R from a peptide library using phage-based cell surface selection is a novel finding, offering a starting point for the selection of novel CB_2_R antagonists which may be used therapeutically in the treatment of IBD for the inhibition of leukocyte trafficking to the gut, especially in a time where the clinical need continues to rise. Given the success in specifically targeting CB_2_R, this approach could be adopted for the targeting of other receptors involved in diseases where there remains an unmet clinical need.

## Methods

### Jurkat T Cell Culture

Jurkat clone E6-1 cells, a human T cell lymphocytic leukaemia cell line, were obtained from the American Type Culture Collection. The cells were maintained in RPMI-1640 media containing L-glutamine supplemented with penicillin-streptomycin and fetal bovine serum (FBS) to a final concentration of 1% and 10%, respectively. Cells were incubated at 37°C (95% O_2_/5% CO_2_). Media was changed and cells were passed every 2-3 days.

### *CNR2* CRISPRa HEK293T Cell Culture

HEK293T cells were seeded in a 6-well plate at a seeding density of 6.25 × 10⁵ cells in a total volume of 2 ml 24 h prior to transfection and allowed to adhere overnight. On the day of transfection, the culture medium was aspirated and replaced with 2 ml of fresh DMEM.

DNA-lipid complexes were prepared in 1.5 ml microcentrifuge tubes for each treatment condition. Each complex was assembled in a total volume of 500 µl Opti-MEM. The transfection conditions were as follows: Well 1 served as an untransfected vehicle control (500 µl Opti-MEM only); Well 2 received 2.1 µg of LRG-GFP plasmid and served as a transfection efficiency positive control; Well 3 received a three-plasmid combination consisting of 0.54 µg PB-SAM, 1.56 µg PB-sgRNA3, and 0.4 µg of PBase transposase. The concentration of each plasmid was determined by NanoDrop spectrophotometry prior to transfection in order to calculate the required volume. PBase (0.5 µg/µl) was added at a volume of 0.8 µl to achieve the desired mass of 0.4 µg.

To each DNA-Opti-MEM mixture, 4 µl of Lipofectamine LTX reagent and 2.5 µl of PLUS reagent were added as per the manufacturer’s recommendations. Complexes were mixed gently by pipetting and incubated at room temperature for 30 minutes. Following incubation, each complex was transferred to its designated well. Cells were incubated for 48 hours post-transfection prior to the initiation of antibiotic selection.

Stable transformants were selected by supplementing the culture medium with Hygromycin B at a concentration of 300 µg/ml, a dose determined empirically by antibiotic kill curve. Selection was maintained for a minimum of 9 days. Once cells reached confluency in the 6-well plate, they were expanded into larger tissue culture flasks. The resulting CRISPRa *CNR2*-HEK293T cell line was subsequently maintained in culture medium supplemented with 300 µg/ml Hygromycin B.

### HTLA Cell Culture

The HEK293-derived cell line containing stable integrations of a tetracycline transactivator-dependent luciferase reporter and a β-arrestin2-tobacco etch virus (TEV) fusion gene (HTLA cells) were obtained from….. The cells were maintained in Dulbecco’s Modified Eagle Medium (DMEM), supplemented with 100 μg/ml hygromycin B, 2 μg/ml puromycin, penicillin-streptomycin and FBS to a final concentration 1%, and 10%, respectively. Cells were incubated at 37 °C (5% CO_2_/95% air). Media was changed and cells were passed every 3-4 days.

### Cell Surface Selection against *CNR2*

#### Day One

A confluent T75 of Jurkat T cells was pooled and pelleted via centrifugation at 1,200 × *g* at 25 °C for 5 minutes prior to aspirating the cell culture media. The cells were rinsed twice in 1X phosphate-buffered saline (PBS) via centrifugation at 1,200 × *g* at 25 °C for 5 minutes before being resuspended 8.5 ml of PBS/0.05% Tween/5% bovine serum albumin (BSA) and 750 μl each of 7mer and 5mer peptides. Resuspended cells were incubated on an over and under rotator at room temperature for 5 minutes, pelleted via centrifugation at 1,200 × *g* at 25 °C for 5 minutes to aspirate unbound phage, and washed three times in 10 ml of PBS/0.05% Tween/5% BSA under the same centrifugation conditions. Washed cells were resuspended in 1 ml of 0.1 M pH 2.2 filter sterilised glycine, transferred to 1.5 ml microcentrifuge tubes, and incubated at room temperature on an over and under rotator for 4 minutes. The cell biomass was pelleted via centrifugation at 1,500 × *g* at 25 °C for 3 minutes and the cleared aspirate was removed and immediately neutralised by adding to 2 ml of 1 M Tris pH 7.4, prior to being stored at 4 °C.

A starter culture of TG1 E. coli cells was prepared by adding a 500 μl stock aliquot to 50 ml of sterilised 2xYT (made up of 8g tryptone, 5 g yeast extract and 2.5 g NaCl in 500 ml deionised H_2_O) media in an autoclaved 250 ml flask under a flame, and was grown in a shaking incubator at 37 °C, 240 rpm. Just before reaching OD_600_ 0.4, the TG1 flask was removed from the incubator and immediately placed in a 37 °C water bath to avoid TG1 cooling. 4.5 ml of the TG1s was removed using a pipette gun and added to 1.5 ml of the previously prepared neutralised, clear phage eluate, prior to being incubated in a 37 °C water bath for 30 minutes. Following 30 minute infection, 100 μl was removed from the tube and 1/100, 1/1,000, and 1/10,000 serial dilutions were prepared using sterilised 2xYT media under a flame. The remaining contents were pelleted via centrifugation at 9,000 × *g* for 2 minutes. The supernatant was decanted and infected TG1s were resuspended in 200 μl of sterilised 2xYT under a flame. This 200 μl was spread on a large 100 mm cell culture dish containing TYE/100 ng/μL ampicillin /1% glucose and stored overnight at 37 °C and will be used as the lawn. 10 μl of the dilution plates were spread on small 10mm cell culture dishes in triplicate containing TYE, 100 ng/μL ampicillin, and 1% glucose and stored overnight at 37 °C.

#### Day Two

Following overnight incubation, dilution plates were used to calculate colony-forming units (CFU)/ml and TG1 colonies were scraped off the lawn in 2 ml of sterilised 2xYT media containing 15% glycerol under a flame. 50 μl of this bacterial stock was added to 50 ml sterilised 2xYT media containing 100 μg/ml ampicillin and 1% glucose in an autoclaved 250 ml flask and placed in a shaking incubator at 37 °C, 240 rpm. Just before reaching OD_600_ 0.4, the TG1 flask was removed from the incubator and immediately placed in a 37 °C water bath to avoid TG1 cooling. 10 ml of this cell culture was then infected with approximately 5×10^10^ KM13 helper phage for efficient phage rescue, and incubated in a 37 °C water bath for 30 minutes. Following incubation, the cells were pelleted via centrifugation at 3,300 × *g* for 10 minutes at 4 °C. and resuspended in 50 ml sterilised 2xYT media containing 100 μg/ml ampicillin, 50 μg/ml kanamycin, and 0.1% glucose in an autoclaved 250 ml flask. This was then grown overnight in a shaking incubator at 30 °C, 220 rpm.

#### Day Three

Overnight culture was pelleted via centrifugation at 3,300 × *g* for 15 minutes at 4 °C. 40 ml of the supernatant was saved and precipitated with 10 ml PEG/NaCl on ice for 1 hour. After an hour, this was pelleted via centrifugation at 3,300 × *g* for 30 minutes at 4 °C, and the pellet was resuspend in 2 ml of PBS/15% glycerol. The resuspension was again pelleted via centrifugation at 11,600 × *g* for 10 minutes at room temperature in a microcentrifuge tube, and the clear supernatant phage stock was labelled as ‘Round 1’ and stored at 4 °C for short term storage or at -80 °C for long term storage.

Round 2 of cell surface selection was completed on *CNR2* CRISPRa HEK293T cells. Three 10cm^3^ plates of *CNR2* CRISPRa HEK293T cells at > 60% confluency were rinsed in pre-warmed 1X PBS before incubation in 5 ml pre-warmed FBS-free DMEM supplemented with 7 mM Ethylenediaminetetraacetic Acid (EDTA) for 5 minutes at 37 °C. The plates were gently knocked alongside some additional pipetting of media to dislodge any remaining adherent cells and the cells were pooled into a 15 ml tube. The cells were pelleted via centrifugation at 1,200 × *g* at 25 °C for 5 minutes prior to aspirating the EDTA-containing media. The cells were rinsed twice in 1X PBS via centrifugation at 1,200 × *g* at 25 °C for 5 minutes before being resuspended 9 ml of PBS/0.05% Tween/5% BSA and 1 ml of the previously prepared Round 1 phage stock. The protocol was then followed as described above.

Round 3 of cell surface selection was completed on wild-type Jurkat T cells. A confluent T75 of Jurkat T cells was pooled and pelleted via centrifugation at 1,200 × *g* at 25 °C for 5 minutes prior to aspirating the cell culture media. The cells were pelleted via centrifugation at 1,200 × *g* at 25 °C for 5 minutes prior to aspirating the cell culture media. The cells were rinsed twice in 1X PBS via centrifugation at 1,200 × *g* at 25 °C for 5 minutes before being resuspended 8.5 ml of PBS/0.05% Tween/5% BSA and 1 ml of the previously prepared Round 2 phage stock. The protocol was then followed as described above.

### Polymerase Chain Reaction (PCR) and Gel Purification of Round 2 and Round 3 Phage Stocks

Phage stocks from Round 2 and Round 3 of cell surface selection were prepared for next generation sequencing (NGS). 500 μl of each phage stock was pelleted via centrifugation at 6,800 × *g* for 3 minutes at room temperature, and the DNA samples were prepared using the QiaPrep^®^ Spin Miniprep Kit according to the manufacturer’s instructions. 5 ng of prepared DNA was PCR amplified in a master mix containing 10 μl 5X Q5^®^ PCR reaction buffer, 1 μl dNTPs, 1.2 μl 500 nM forward and reverse S100G primers, and 0.5 μl Q5^®^ high-fidelity DNA polymerase (New England Biolabs; M0491S) made up to a final reaction volume of 50 μl using sterilised de-ionised H_2_O. These samples were placed in a thermocycler and ran at the following conditions for efficient PCR amplification of prepared DNA:

**Table 1:** Thermocycler conditions for PCR amplification.

| Temperature (°C) | Time (s) |  |
| --- | --- | --- |
| 98°C | 30s | } x 1 |
| 98°C | 10s |  |
| 55°C | 20s | } x 25 |
| 72°C | 10s |  |
| 72°C | 120s |  |
| 4°C | 300s |  |

40 μl of prepared PCR product was mixed with 12 μl of 6X DNA loading dye, 50 μl of which was loaded into a 2% agarose gel made with 1.4 g agarose, 70 ml 1X TAE, and 7 μl SYBR safe DNA gel stain. 10 μl of 100 base pair DNA ladder was used as the loading control. The gel was initially run at 65 V for 15 minutes, before being increased to between 80-90 V for 1-2 h. Following sufficient run time, the gel was visualised using the Vilber E Box, and the respective DNA bands were excised accordingly. DNA from the excised gel was then efficiently extracted and cleaned up using the QiaQuick^®^ gel extraction kit according to the manufacturer’s instructions. The concentration of the extracted DNA samples was calculated using the NanoDrop, and the samples were sent to NeoChromosome Inc. for NGS.

### BL21 Star Competent Cell Transformation

On ice, one vial of competent OneShot^™^ BL21 Star^™^ E. coli cells was thawed per each required transformation. The required pET-3a vector plasmid containing the respective CB_2_R-specific sequence cloned in via Nde/BamHI cloning sites (GenScript) was vortexed briefly and placed on ice. From the plasmid, 1 µl was added to 17 μl of competent cell stock and mixed gently by tapping, before being incubated on ice for 30 minutes. The remaining plasmid was stored at - 20°C. Following incubation the E. coli cells were heat shocked using a heat block for 45 seconds at 42 °C in the absence of any shaking. Immediately following the heat shock period, the vial was removed and placed on ice for 5 minutes. The transformed cells were then plated on pre-warmed selective petri dish containing LB and agar, supplemented with 100 µg/ml ampicillin and chloramphenicol and incubated at 37 °C overnight for selective growth.

### Bioinformatics Pipeline for NGS Data Analysis

The results of NGS from NeoChromosome Inc. were obtained in the FASTQ format along with sequence quality report. A parser step was completed to break FASTQ files into CSV files with each row containing sequence index, FASTQ sequence and associated quality scores. These CSV were then read by the Rev_For_Search module, which scanned for forward and reverse adapter sequences, as well as linkers flanking the NNK-format insert. This categorised the sequences into forward reads, reverse-complemented sequences, and orphan sequences lacking identifiable primers which were filtered out from the input. The Translate_Clean step took the mapped nucleotide sequences, evaluated the quality of the NNK regions, and translated them into amino acid sequences. To determine the correct reading frame, the script scanned all three frames and identified the one containing 7x [NNK] flanked by amino acid tags. Next, the Frequency Count module compiled the translated the 7-mer peptide inserts, calculated the frequency of each unique peptide, and summarised the distribution within the screen library. Finally, the Clustering step applied an amino acid scoring matrix, as developed by Linse *et al.* [17] to cluster peptides by aligning each sequence against all others in the dataset, with a chosen cutoff alignment score of ≥ 30.

### Microscale Expression and Purification of CB_2_R-specific Peptides for Functional Analysis

Single isolated colonies from selection plates were picked and inoculate into 10 ml of Overnight Express^™^ Instant TB media supplemented with filter sterilised 1% glycerol 100 μg/ml ampicillin. The inoculated samples were incubated in a shaking incubator at 37 °C for 20-24 h for efficient protein expression. Following incubation, the samples were pelleted via centrifugation at full speed (4500-5000 × g) at 4 °C for 15 minutes. The supernatant was discarded carefully and the pellets were resuspended in 1 ml of 10 mM Tris pH 8.0 and transferred to 1.5 ml microcentrifuge tubes, before being placed in a sonicating water bath for 10 minutes. The resulting cell lysate was frozen at -20 °C for 2 h and then thawed in a water bath at 37 °C for 10 minutes prior to boiling. The lids of respective microcentrifuge tubes were pierced carefully with a syringe needle, and the tubes were placed in a heat block and boiled at 100 °C for 10 minutes. Following boiling, the tubes were removed carefully and pelleted via centrifugation at 18,000 × *g* for 15 minutes. The resulting supernatant were saved, added to 5 μl of DNase in new microcentrifuge tubes and placed in a heat block at 37 °C for 30 minutes. Following incubation, 15 μl of 200 mM EDTA stock was added to each tube for a final EDTA concentration of 3 mM, prior to being added to 9 ml of 10 mM Tris pH 8.0/1 mM EDTA and purified via ion exchange chromatography (IEX) for downstream functional analysis. Add 100 μl of DEAE Sephacel^™^ resin to each well of an AcroPrep^™^ Advance 96-well 10-30 K filtration plate. Wash the resin multiple times with 10 mM 4-(2-Hydroxyethyl)-1-piperazineethanesulfonic Acid (HEPES)/Tris and 1 mM EDTA pH 8.0 in order to achieve a resin pH of 8.0. 200 μl of protein eluate was added to respective wells and leave rest for 10 minutes. The plate was spun at room temperature for 5 minutes at 2,000 × *g* and the resulting flow through in the collection plate was transferred to fresh microcentrifuge tubes. These elution steps were repeated with 200 μl 10 mM HEPES/Tris and 1 mM EDTA pH 8.0 + a 0-500 mM serial dilution of NaCl. The flow through from each elution was collected and a sodium dodecyl sulfate-polyacrylamide gel electrophoresis (SDS-PAGE) gel was run to confirm the presence of each sample prior to the second IEX. The eluates were then pooled and another SDS-PAGE gel was run to confirm eluates with 10 kDa bands as S100G selektides. The protein concentration A280 of each pooled sample was measured using the NanoDrop, and based on the resin binding capacity the pooled samples were diluted with 10 mM HEPES/Tris and 2 mM CaCl_2_ or water. In a fresh 96-well filtration plate, 800 μl of resin was added to each well and multiple washes were completed as above. Following sufficient washing, 1.5 ml of diluted sample was added to the resin and left sit for 5 minutes. The plate was spun at room temperature for 5 minutes at 2,000 × *g* and the resulting flow through in the collection plate was collected and transferred to a fresh 15 ml tube. This was repeated until all of the sample was loaded on to the resin. These elution steps were repeated with 800 μl 10 mM HEPES/Tris and 2 mM CaCl_2_ + a 0-300 mM serial dilution of NaCl. The flow through from each elution was collected and an SDS-PAGE gel was run to confirm the presence of each sample. The eluates were then pooled and another SDS-PAGE gel was run to confirm eluates with 10 kDa bands as S100G selektides. The protein concentration A280 of each pooled sample was measured using the NanoDrop, being diluted accordingly for future use in functional assays.

### Fluorescent Glucose Analogue ptake Assay in Jurkat T cells

In 100 µl of pre-warmed supplemented RPMI-1640, Jurkat T cells were plated at approximately 1×10^6^ cells/ml into a clear, sterile, flat-bottom 96-well plate. For 3 h and in biological and technical triplicates, the cells were exposed simultaneously to 1 μM CB_2_R agonist JWH-133 (JWH) alongside varying concentrations of the purified selektides freshly prepared in glucose-free RPMI-1640 media, in order to assess the effect of the selektides on CB_2_R-driven glucose uptake in Jurkat T cells. Vehicle-treated cells received 100% dimethyl sulfoxide (DMSO) diluted in the same manner as JWH-133 in glucose-free media in the place of any cannabinoid treatment. 10 minutes prior to treatment time concluding the cells were exposed to the fluorescent glucose analogue 2-(N-(7-nitrobenz-2-oxa-1,3-diazol-4-yl)amino)-2-deoxyglucose (2-NBDG) prepared in glucose-free media, at a final concentration of 100 µM in each well. The cells were again incubated at 37 °C for 10 minutes. Following 10 minute incubation, the plate was removed from the incubator and the cells were transferred to a white opaque, flat-bottom 96 well plate. The cells were then pelleted by centrifugation at 400 × *g* at room temperature for 5 minutes using the Eppendorf Centrifuge 5810R. Without disturbing the cell layer, the supernatant was removed, before 200 µl of cell-based assay buffer/PBS was added to each well to sufficiently wash the cells. Again, the plate was centrifuged for 5 minutes at 400 × *g* at room temperature. Similarly, the supernatant was removed carefully as to not disturb the cell layer, before 100 µl of cell-based assay buffer/PBS was added to each well. The fluorescent intensity of the treated cells was then analysed immediately using the ClarioStar PLUS plate reader (BMG LabTech) with filters designed to detect fluorescence at Ex/Em of 485nm/535nm.

### Phosphorylated ERK Level Assessment in Jurkat T cells

In 100 μl of pre-warmed supplemented RPMI-1640, Jurkat T cells were plated at approximately 2 x10^6^ cells/ml into a clear V-bottom 24-well plate. In biological and technical triplicates, the cells were simultaneously exposed to 1 μM CB_2_R agonist JWH-133 alongside 100 nM of selektides for 30 minutes, freshly prepared in glucose-free RPMI-1640 media, in order to assess the effect of the selektides on downstream intracellular ERK1/2 signalling following CB_2_R activation. Vehicle-treated cells received 100% DMSO diluted in the same manner as JWH-133 in RPMI-1640 media in the place of any cannabinoid/Selektide treatment. Following treatment, the cells were pelleted via centrifugation for 5 minutes at 1500 × *g* using the Eppendorf centrifuge 5810R. Pelleted cells were then intracellularly stained with anti-human/mouse phospho-ERK1/2 T202/Y204 APC following fixation and permeabilisation using the Intracellular Fixation and Permeabilisation Buffer Set according to the manufacturer’s’ instructions. Live cells were identified using Live/Dead Fixable Aqua dye. Following respective fixation and permeabilisation, the cells were resuspended in 100 μl PBS, before being analysed using the Cytoflex LX system (Beckman Coulter), and post-run analyses were performed using CytExpert software (Beckman Coulter).

### β-arrestin Recruitment Assessment in HTLA cells

#### Day One

In 100 μl of pre-warmed supplemented DMEM, HTLA cells were plated at approximately 0.5 x10^6^ in a sterile, flat-bottom 96 well plate.

#### Day Two

100 ng CB_2_R-Tango DNA was transfected per well using Lipofectamine 2000 according to the manufacturer’s instructions. Non-transfected control wells received 10 μl Opti-MEM reduced serum media in place of any DNA complex.

#### Day Three

Following transfection, the cellular media and transfection mix were removed from the cells prior to treatment and the cells were washed once in 1X PBS. The cells were simultaneously exposed to varying concentrations of JWH-133 alongside a fixed concentration of selektides at 100 nM for 24 h, freshly prepared in supplemented DMEM media. Vehicle-treated cells received 100% DMSO diluted in the same manner as JWH-133 in DMEM media in the place of any cannabinoid/Selektide treatment.

#### Day Four

After 24 h stimulation, the cells were removed from incubation and 50 μl of the Promega Bright-Glo Luciferase Assay System was added to each well according to the manufacturer’s instructions. The plate was incubated in the dark for 20 minutes at room temperature before the luminescent signal was measured using the ClarioStar PLUS plate reader.

### Mice Handling

TNF^ΔARE/+^ mice (B6.129S-Tnftm2GKI/Jarn; MGI:3720980) were generated by backcrossing heterozygous TNF^ΔARE/+^ mice to C57BL/6J. Male and female TNF^ΔARE/+^ mice aged 8–12 weeks were randomised across the three tested treatment groups. Animals were maintained under specific pathogen-free conditions with *ad libitum* access to standard chow and water. All experimental procedures were approved by the Institutional Animal Care and Use Committee of the University of Colorado.

### Single-Cell Suspension Preparation and Flow Cytometry

Single-cell suspensions were obtained from the spleen and mesenteric lymph nodes (MLN) by mechanical dissociation through a 70 µm cell strainer. Erythrocytes were depleted from splenocyte preparations by incubation in ammonium chloride lysing reagent. Lamina propria mononuclear cells (LPMCs) were isolated as previously described. Briefly, intestinal tissues were agitated in saline solution containing 1 mM EDTA to liberate epithelial cells, followed by digestion of the remaining stromal tissue with 1 mg/ml Collagenase VII (Sigma-Aldrich: C0773). Single-cell suspensions from each tissue compartment were washed in PBS and stained with the relevant fluorophore-conjugated antibody combinations for surface marker detection. Flow cytometry data were acquired using the BD FACSCanto II and analysed using FCS Express.

### Statistics

Statistical analyses were performed using Student t test. Graphs are presented as means ± standard error of the mean (SEM) and were generated using GraphPad Prism 10 software. Values of p < 0.05 were considered statistically significant.

## Notes

### Competing Interest Statement

The authors have declared no competing interest.

### Funding

- Enterprise Ireland: CF20242332

## Bibliography

1. Mowat, A.M. and W.W. Agace, Regional specialization within the intestinal immune system. Nature reviews. Immunology, 2014. 14(10): p. 667–685.

2. Collins, C.B., et al., Flt3 ligand expands CD103+ dendritic cells and FoxP3+ T regulatory cells, and attenuates Crohn’s-like murine ileitis. Gut, 2012. 61(8): p. 1154–1162.

3. Ng, S.C., et al., Worldwide incidence and prevalence of inflammatory bowel disease in the 21st century: a systematic review of population-based studies. The Lancet (British edition), 2017. 390(10114): p. 2769–2778.

4. Kaplan, G.G. and J.W. Windsor, The four epidemiological stages in the global evolution of inflammatory bowel disease. Nature reviews. Gastroenterology & hepatology, 2021. 18(1): p. 56–66.

5. Kumar, A., et al., Crossing barriers: the burden of inflammatory bowel disease across Western Europe. Therapeutic Advances in Gastroenterology, 2023. 16: p. 17562848231218615.

6. Sebastian, S., et al., Promoting equity in inflammatory bowel disease: a global approach to care. The Lancet Gastroenterology & Hepatology, 2024. 9(3): p. 192–194.

7. Yanai, H. and S.B. Hanauer, Assessing response and loss of response to biological therapies in IBD. Official journal of the American College of Gastroenterology| ACG, 2011. 106(4): p. 685–698.

8. Feagan, B.G., et al., Vedolizumab as Induction and Maintenance Therapy for Ulcerative Colitis. The New England journal of medicine, 2013. 369(8): p. 699–710.

9. Sandborn, W.J., et al., Vedolizumab as Induction and Maintenance Therapy for Crohn’s Disease. The New England journal of medicine, 2013. 369(8): p. 711–721.

10. Eriksson, C., et al., Real-world effectiveness of vedolizumab in inflammatory bowel disease: week 52 results from the Swedish prospective multicentre SVEAH study. Therapeutic advances in gastroenterology, 2021. 14: p. 17562848211023386.

11. Ambrose, T. and A. Simmons, Cannabis, cannabinoids, and the endocannabinoid system—is there therapeutic potential for inflammatory bowel disease? Journal of Crohn’s and Colitis, 2019. 13(4): p. 525–535.

12. Koning, M., et al., Use and predictors of oral complementary and alternative medicine by patients with inflammatory bowel disease: a population-based, case-control study. Inflamm Bowel Dis, 2013. 19(4): p. 767–78.

13. Hasenoehrl, C., M. Storr, and R. Schicho, Cannabinoids for treating inflammatory bowel diseases: where are we and where do we go? Expert review of gastroenterology & hepatology, 2017. 11(4): p. 329–337.

14. Galiègue, S., et al., Expression of central and peripheral cannabinoid receptors in human immune tissues and leukocyte subpopulations. European journal of biochemistry, 1995. 232(1): p. 54–61.

15. Schatz, A.R., et al., Cannabinoid Receptors CB1 and CB2: A Characterization of Expression and Adenylate Cyclase Modulation within the Immune System. Toxicology and applied pharmacology, 1997. 142(2): p. 278–287.

16. Leddy, R.S., et al., CB2R promotes T cell gut homing and exacerbates ileitis in a murine Crohn’s model. Inflammatory bowel diseases, 2026.

17. Linse, S., et al., An aggregation inhibitor specific to oligomeric intermediates of Aβ42 derived from phage display libraries of stable, small proteins. Proceedings of the National Academy of Sciences - PNAS, 2022. 119(21): p. 1–10.

18. Xu, H., et al., Anti-inflammatory property of the cannabinoid receptor-2-selective agonist JWH-133 in a rodent model of autoimmune uveoretinitis. Journal of Leucocyte Biology, 2007. 82(3): p. 532–541.

19. Montecucco, F., et al., CB2 cannabinoid receptor activation is cardioprotective in a mouse model of ischemia/reperfusion. Journal of molecular and cellular cardiology, 2009. 46(5): p. 612–620.

20. Weiss, A. and F. Friedenberg, Patterns of cannabis use in patients with Inflammatory Bowel Disease: A population based analysis. Drug and alcohol dependence, 2015. 156: p. 84–89.

21. Phatak, U.P., et al., Prevalence and Patterns of Marijuana Use in Young Adults With Inflammatory Bowel Disease. Journal of pediatric gastroenterology and nutrition, 2017. 64(2): p. 261–264.

22. Naftali, T., et al., Cannabis Induces a Clinical Response in Patients With Crohn’s Disease: A Prospective Placebo-Controlled Study. Clinical gastroenterology and hepatology, 2013. 11(10): p. 1276–1280.e1.

23. Storr, M., et al., Cannabis use provides symptom relief in patients with inflammatory bowel disease but is associated with worse disease prognosis in patients with Crohn’s disease. Inflammatory bowel diseases, 2014. 20(3): p. 472–480.

24. Leinwand, K.L., et al., Cannabinoid receptor-2 ameliorates inflammation in murine model of Crohn’s disease. Journal of Crohn’s and Colitis, 2017. 11(11): p. 1369–1380.

25. Storr, M.A., et al., Activation of the cannabinoid 2 receptor (CB2) protects against experimental colitis. Inflammatory bowel diseases, 2009. 15(11): p. 1678–1685.

26. Stintzing, S., et al., Role of cannabinoid receptors and RAGE in inflammatory bowel disease. Histology and histopathology, Vol. 26, n°6 (2011), 2011.

27. Strisciuglio, C., et al., Increased expression of CB2 receptor in the intestinal biopsies of children with inflammatory bowel disease. Pediatric research, 2022.

28. Wright, K., et al., Differential expression of cannabinoid receptors in the human colon: cannabinoids promote epithelial wound healing. Gastroenterology, 2005. 129(2): p. 437–53.

29. Hryhorowicz, S., et al., Endocannabinoid system as a promising therapeutic target in inflammatory bowel disease–a systematic review. Frontiers in immunology, 2021. 12: p. 790803.

30. Coopman, K., et al., Temporal variation in CB2R levels following T lymphocyte activation: Evidence that cannabinoids modulate CXCL12-induced chemotaxis. International immunopharmacology, 2007. 7(3): p. 360–371.

31. Marquéz, L., et al., Ulcerative colitis induces changes on the expression of the endocannabinoid system in the human colonic tissue. PloS one, 2009. 4(9): p. e6893–e6893.

32. Ettaro, R., et al., Behavioral assessment of rimonabant under acute and chronic conditions. Behavioural brain research, 2020. 390: p. 112697–112697.

33. Moreira, F.A. and J.A.S. Crippa, The psychiatric side-effects of rimonabant. Revista brasileira de psiquiatria, 2009. 31(2): p. 145–153.

34. Huang, Z.-b., et al., Protective effects of specific cannabinoid receptor 2 agonist GW405833 on concanavalin A-induced acute liver injury in mice. Acta Pharmacologica Sinica, 2019. 40(11): p. 1404–1411.

35. Börner, C., V. Höllt, and J. Kraus, Activation of human T cells induces upregulation of cannabinoid receptor type 1 transcription. Neuroimmunomodulation, 2008. 14(6): p. 281–286.

36. Barnea, G., et al., The genetic design of signaling cascades to record receptor activation. Proceedings of the National Academy of Sciences - PNAS, 2008. 105(1): p. 64–69.

37. Zeghal, M., G. Laroche, and P.M. Giguère, Parallel Interrogation of β-Arrestin2 Recruitment for Ligand Screening on a GPCR-Wide Scale using PRESTO-Tango Assay. Journal of visualized experiments, 2020. 2020(157).

38. Violin, J.D., D.G. Soergel, and M.W. Lark, Beta-arrestin-biased ligands at the AT1R: a novel approach to the treatment of acute heart failure. Drug discovery today. Therapeutic strategies, 2012. 9(4): p. e149–e154.

39. Gesty-Palmer, D., et al., Distinct β-Arrestin- and G Protein-dependent Pathways for Parathyroid Hormone Receptor-stimulated ERK1/2 Activation. The Journal of biological chemistry, 2006. 281(16): p. 10856–10864.

40. Lorente, J.S., et al., GPCR drug discovery: new agents, targets and indications. Nature reviews. Drug discovery, 2025. 24(6): p. 458–479.

41. Riquelme-Sandoval, A., et al., New Insights Into Peptide Cannabinoids: Structure, Biosynthesis and Signaling. Frontiers in pharmacology, 2020. 11: p. 596572.

42. Tomašević, N., et al., Discovery and development of macrocyclic peptide modulators of the cannabinoid 2 receptor. The Journal of biological chemistry, 2024. 300(6): p. 107330.

43. Rossino, G., et al., Peptides as Therapeutic Agents: Challenges and Opportunities in the Green Transition Era. Molecules, 2023. 28(20): p. 7165.

44. Khezri, H., et al., Peptibodies: Bridging the gap between peptides and antibodies. International journal of biological macromolecules, 2024. 278(Pt 2): p. 134718.

45. Roos, A., et al., Specific Inhibition of the Classical Complement Pathway by C1q-Binding Peptides. The Journal of immunology (1950), 2001. 167(12): p. 7052–7059.

46. Fang, Y., et al., Design of p53-derived peptides with cytotoxicity on breast cancer. Amino Acids, 2014. 46(8): p. 2015–2024.

47. Wu, T., et al., Quantification of epitope abundance reveals the effect of direct and cross-presentation on influenza CTL responses. Nature communications, 2019. 10(1): p. 2846–14.

