## Supplementary material for "Selection of potent biologic antagonists of the cannabinoid GPCR CB_2_R from a constrained peptide library": supplemntal figure 1 and 2

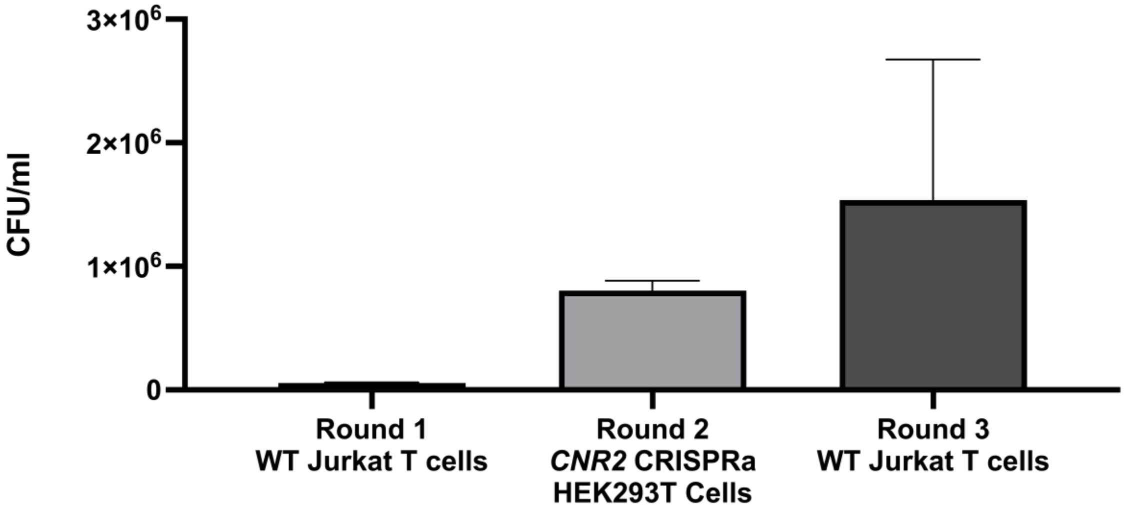


| Selection Round | Cell Type | CFU/ml |
|---|---|---|
| Round 1 | WT Jurkats | 56,000 CFU/ml |
| Round 2 | CNR2 CRISPRa HEKs | 800,000 CFU/ml |
| Round 3 | WT Jurkats | 1.53 million CFU/ml |

**Supplementary Figure S1: Selection outline utilised for cell surface selection against *CNR2* uses endogenously-expressing and genetically modified overexpressing cell lines for efficient peptide binder amplification.** Bar chart and table depicting the CFU/ml values as selection rounds alternate between endogenously *CNR2*-expressing WT Jurkat T cells and genetically modified *CNR2* overexpressing HEK293T cells. (51 words)

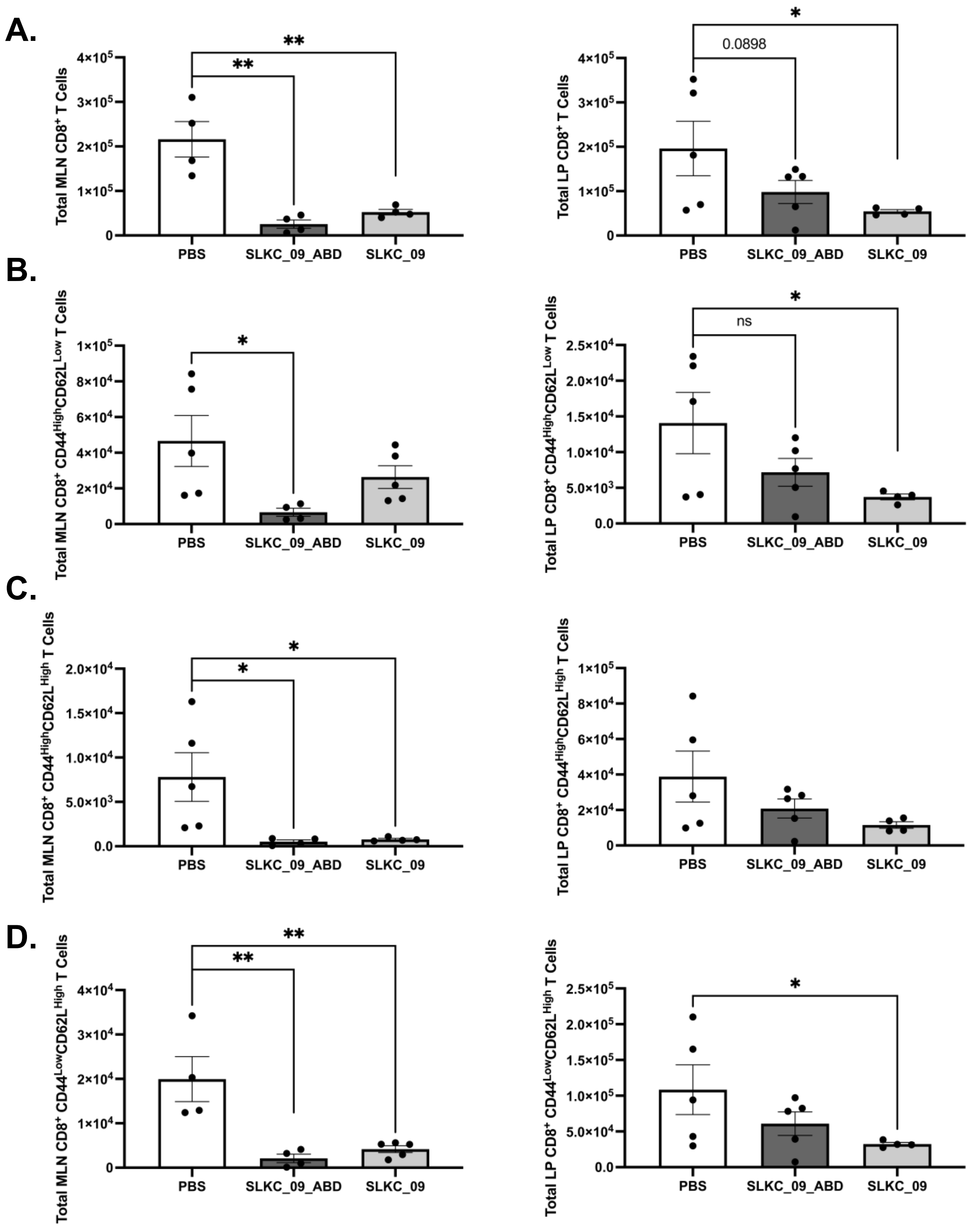


**Supplementary Figure S2: SLKC_09 and SLKC_09_ABD broadly suppress CD8$^+$ T cell subsets across gut-associated lymphoid compartments.** Total CD8$^+$ T cell numbers were quantified by flow cytometry across four phenotypically defined subpopulations in the mesenteric lymph node (MLN) and intestinal lamina propria (LP) following treatment of TNF$^{\Delta ARE}$ mice with SLKC_09_ABD or SLKC_09 relative to PBS vehicle control. **(A)** Total CD8$^+$ T cells in the MLN and LP. **(B)** Effector CD8$^+$CD44$^{High}$CD62L$^{Low}$ T cells in the MLN and LP. **(C)** Central memory CD8$^+$CD44$^{High}$CD62L$^{High}$ T cells in the MLN and LP. **(D)** Naïve

$CD8^+CD44^{Low}CD62L^{High}$ T cells in the LP and MLN. Bars represent mean ± SEM of n = 5 animals per group. Statistical comparisons were made relative to PBS control. * $p < 0.05$; ** $p < 0.01$; p values shown where differences trended toward significance. (133 words)
